# Gestationally-Dependent Immune Organization at the Maternal-Fetal Interface

**DOI:** 10.1101/2021.06.25.449807

**Authors:** Amber R. Moore, Nora Vivanco Gonzalez, Katherine A. Plummer, Olivia R. Mitchel, Harleen Kaur, Moises Rivera, Brian Collica, Theo D. Palmer, Sean C. Bendall

## Abstract

The immune system and placenta have a dynamic relationship across gestation to accommodate fetal growth and development. High-resolution characterization of this maternal-fetal interface is necessary to better understand the immunology of pregnancy and its complications. We developed a single-cell framework to simultaneously immuno-phenotype circulating, endovascular, and tissue-resident cells at the maternal-fetal interface throughout gestation, discriminating maternal and fetal contributions. Our data reveal distinct immune profiles across the endovascular and tissue compartments with tractable dynamics throughout gestation that respond to a systemic immune challenge in a gestationally-dependent manner. We uncover that mononuclear phagocytes and neutrophils drive the temporal immune composition of the placenta with remarkably diverse populations, including PD-L1-expressing subsets having compartmental and early gestational bias. Our approach and accompanying datasets provide a resource for additional investigations into gestational immunology and evoke a more significant role for the innate immune system in establishing the microenvironment of early pregnancy.

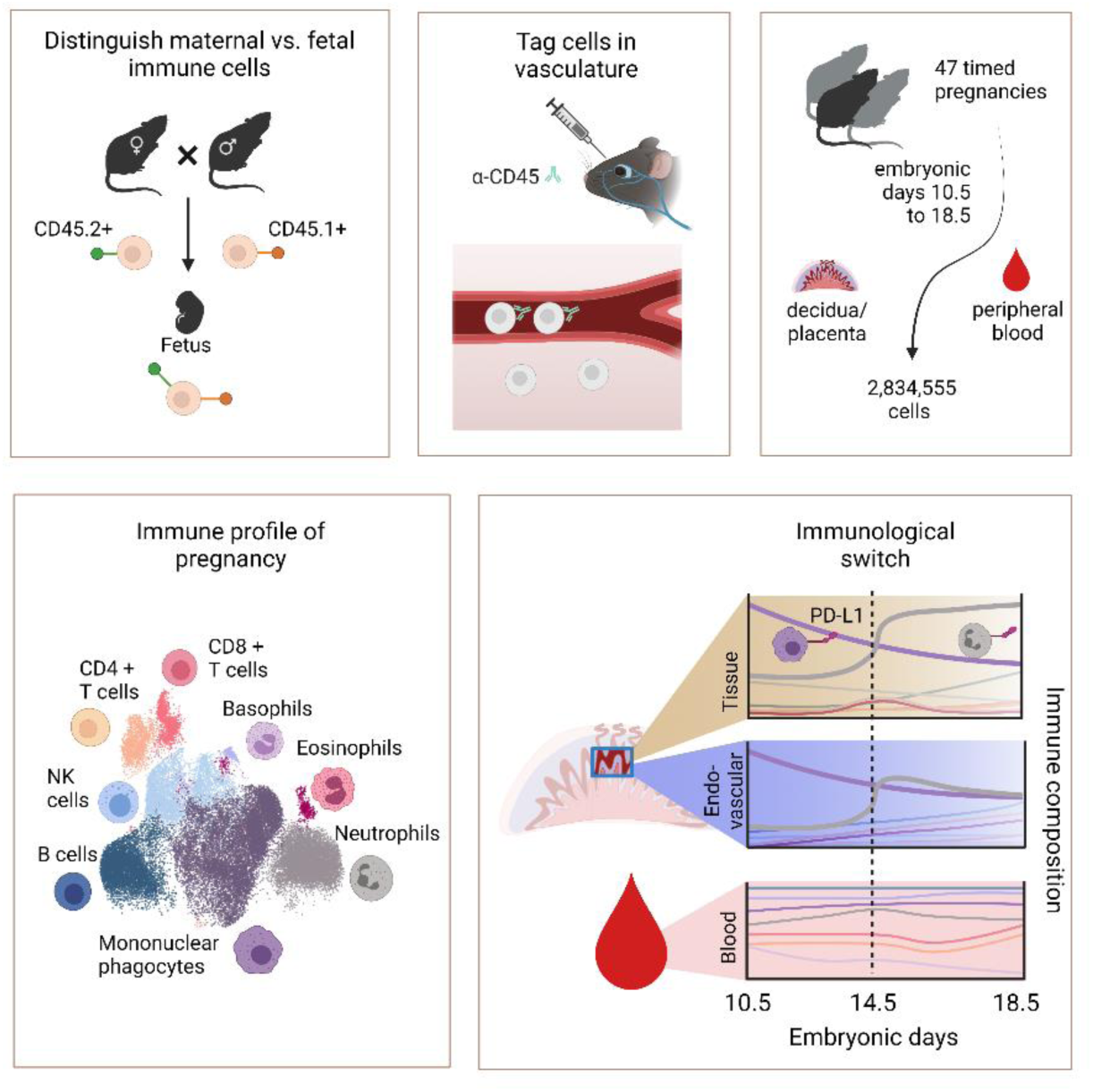

## INTRODUCTION

Though transient, the placenta is a critical multifunctional organ. It mediates nutrient, gas, and waste exchange while simultaneously regulating immune-cell behavior to support tissue remodeling and to maintain tolerance (Ander et al., 2019). The placenta is the site of the maternal-fetal interface, where the fetal chorion is anchored to modified maternal endometrium called the decidua, and where fetal trophoblasts are in direct contact with maternal blood (Ander et al., 2019; Hemberger et al., 2020). Immune regulation at the maternal-fetal interface is critical for pregnancy and healthy fetal development (Ander et al., 2019; Erlebacher, 2013; PrabhuDas et al., 2015). Despite this, detailed knowledge about the dynamics of immune-cell composition and regulation across pregnancy is sparse. A better understanding of the cellular dynamics and homeostasis is necessary for recognizing aberrant immune states, uncovering pathogenic mechanisms, and identifying therapeutic interventions.

Immune-cell phenotypes and function depend on their cellular interactions, tissue localization, and microenvironment compartmentalization (Azizi et al., 2018; Schumacher et al., 2018). The maternal-fetal interface and tumor microenvironment share a number of similarities. Like an invasive tumor, placental architecture is complex and dynamic, requiring cell proliferation, tissue invasion, angiogenesis, vascular remodeling, and modulating tolerance to establish a healthy, functioning maternal-fetal interface (Holtan et al., 2009a; Lala et al., 2021a). As with tumor cell biology, the immune system plays an important role in all of these processes, however, paradoxically, the adaptive arm of the immune system is seemingly dispensable for mammalian pregnancy (Burke et al., 2011; Guleria et al., 2005) implicating a significant role for innate immunity and passive mechanisms for the placentation of a semi-allogeneic fetus. The immune cells both facilitate and respond to this highly dynamic environment, resulting in much adaptation in response to ever changing needs throughout gestation.

Considerable progress has been made in profiling the immune composition of the placenta. Numerous ‘bulk’ transcriptomic approaches have been published and a recent meta-analysis highlights low resolution signatures of pregnancy complications (Yong and Chan, 2020). Flow cytometry and imaging studies shed light on the relative abundance of selected cell types and cytokines (Arenas-Hernandez et al., 2015; Habbeddine et al., 2014; Kruse et al., 1999, 2002; Li et al., 2018; Rowe et al., 2012), and single-cell transcriptomics have begun to resolve many more cell types in first trimester human placentas. For example, decidual natural killer (NK) cells have been resolved into three distinct states and single-cell RNA-seq has predicted potential receptor-ligand interactions that would regulate immune-cell behavior (Vento-Tormo et al., 2018). In addition, an analysis of endovascular and extravascular immune cells in gestational tissues has begun to identify macrophage and dendritic cell signaling related to decidual development (Tagliani et al., 2011). Although the prior studies each contribute to our emerging understanding of pregnancy, the existing data still lacks the depth of analysis necessary to detect rare maternal and fetal immune-cell types or resolve complex populations and their activation states at the single-cell level over time.

We developed a gestational immune monitoring platform to interrogate immune cells at the maternal-fetal interface throughout gestation using a high-dimensional single-cell mass cytometry approach quantifying 40 surface and intracellular markers in conjunction with an unbiased analysis. By crossing congenic mouse strains and applying an injectable antibody to mark circulating cells, we were able to differentiate between maternal and fetal immune cells, as well as endovascular and tissue-resident cells. Applying this to pregnant mice from gestational days E10.5 to E18.5, we were able to detect an immunological switch point that coincides with a molecular switch point and immune reorganization driven by mononuclear phagocytes and neutrophils, many of which bare regulatory proteins like PD-L1. These early and late phases are marked by fluxes in diverse mononuclear phagocyte and neutrophil phenotypes in a compartment-specific manner. With this as a baseline, we applied an in vivo immune perturbation to pregnancy by administering the viral mimic Poly(I:C) on different gestational days. Here, we show that the systemic immune perturbation results in a reduction of PD-L1- expressing mononuclear phagocytes and neutrophils that is both gestational age-dependent and compartment-specific.

We collected 2,834,555 cells with maternal and fetal origin, across two organs, from forty-seven mice over the course of nine gestational days (E10.5-E18.5). We have organized this dataset into an accessible resource for dynamic immune-cell composition at the maternal-fetal interface that is easily mined for future experimental considerations. We further provide a framework to study and interrogate systemic immune perturbation in pregnancy that should have broad adaptability in future studies. Overall, our study reveals a surprisingly dynamic phenotypic diversity that low-dimensional methods and traditionally biased analyses have left concealed and should enable future cross-gestational and inter-compartmental interrogations of the immune systems’ role in pregnancy.

## RESULTS

### A distinct immune composition exists between placental endovascular and peripheral blood

Deep characterization of the immune content at the maternal-fetal interface is needed to elucidate both tissue homeostasis and immune tolerance at higher resolution during pregnancy. We employed single-cell mass cytometry to map maternal immune-cell populations at the maternal-fetal interface during the second half of mouse gestation. Both mice and humans have a hemochorial placenta and share a similar decidual immune composition (Hemberger et al., 2020; Woods et al., 2018). We designed a mass cytometry antibody panel (Table S1) including 37 surface and 3 intracellular markers to identify the immune phenotypes present. We collected placentas with intact maternal decidua from pregnant C57BL/6 dams from embryonic days 10.5 to 18.5. Our ability to differentiate between maternal and fetal immune cells relied on a mating strategy crossing CD45.2 females and CD45.1 males, which produces CD45.2+CD45.1+ fetal immune cells (Figure S1A). To partition immune cells in the placenta by their tissue and endovascular localization, pregnant mice were retro-orbitally injected with an anti-CD45 antibody (Figures 1A, S1F, and S1G), which labels cells in the endovascular space but not parenchymal cells (Anderson et al., 2014; Tagliani et al., 2011). To characterize immune cells using an unbiased approach, mass cytometry data were visualized using the UMAP dimension reduction technique and cells were clustered using the Leiden algorithm. Data pooled from all embryonic days and compartments defined discrete clusters for neutrophils, eosinophils, basophils, NK cells, B cells, T cells, and mononuclear phagocytes (i.e., monocytes, dendritic cells, and macrophages) (Figures 1B and S1B). Cell clusters were metaclustered and classified based on their expression of common immune lineage markers (Figures 1C, 1D, and S1C).

**Figure 1.**
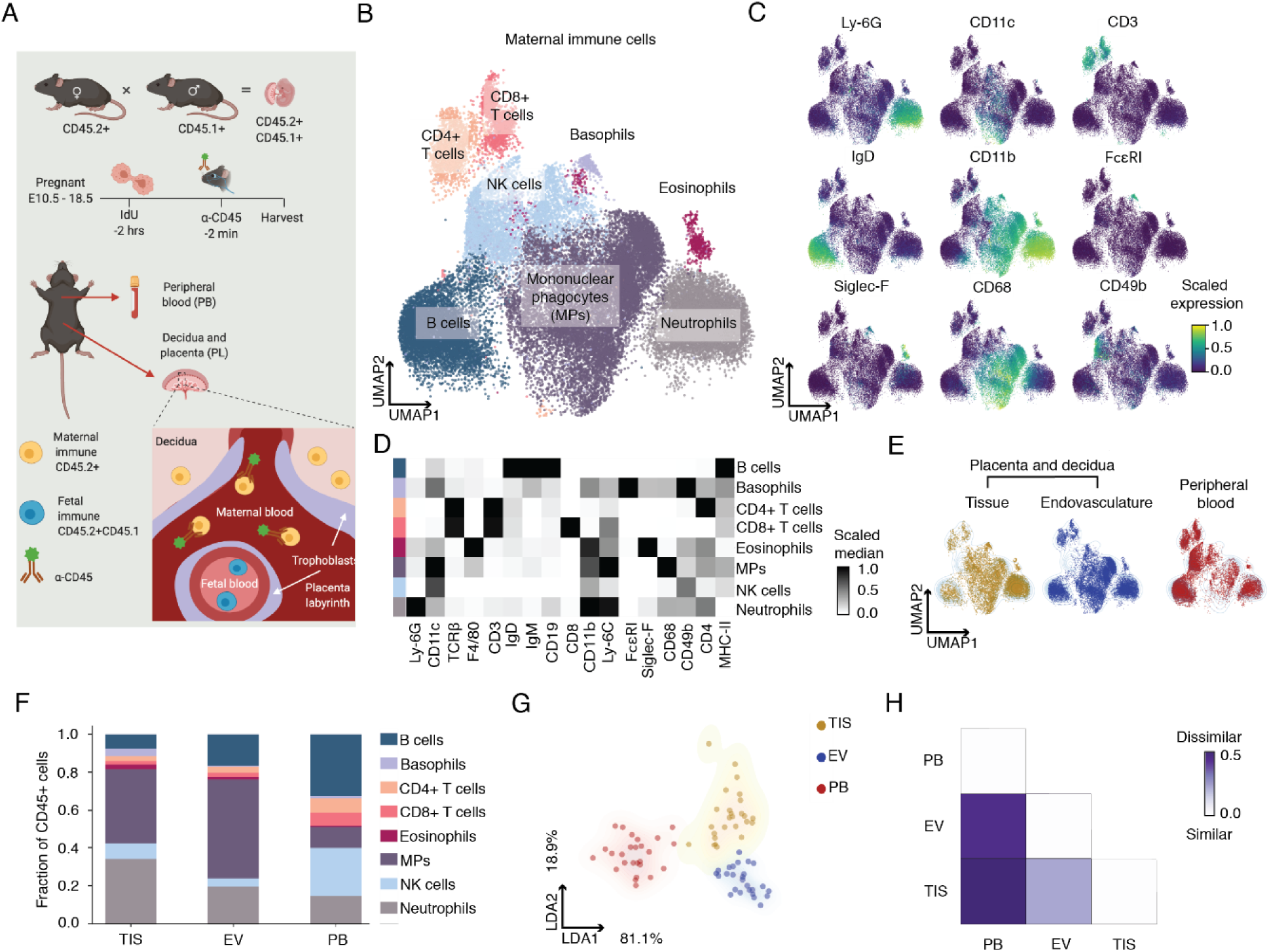
Single-cell mass cytometry reveals distinct immune composition between placental endovasculature and peripheral blood. (A) Experimental set up distinguishes between maternal and fetal immune cells and enables detection of immune cells in the vascular region of the placenta. (B) Organs from 26 mice were used to generate a composite UMAP graph of maternal immune cells in placenta and peripheral blood across E10.5 to 18.5 (C) Scaled cellular median intensity of immune-lineage markers. (D) Scaled median expression of protein markers used for Leiden clustering across maternal immune-cell types identified. (E) Distribution of maternal immune cells across decidua and placenta tissue (TIS) and endovascular (EV) and maternal peripheral blood (PB) compartments projected onto composite UMAP graph as contour plot. (F) Fraction of immune cells relative to total in each compartment. Embryonic days were aggregated. (G) Linear discriminant analysis (LDA) based on maternal immune cell fractions in each compartment. Each dot represents one mouse sample. (H) Bray-Curtis dissimilarity based on maternal immune cell fractions in each compartment.

Tissue microenvironments are known to regulate immune-cell access and function. To examine the impact of placental microenvironments on immune-cell profiles, UMAP projections of composite data were overlaid with immune cells from maternal peripheral blood (PB), immune cells from the endovascular space of the placenta (EV), or immune cells from within the placental tissue (TIS) (Figure 1E). We hypothesized that because the maternal peripheral blood perfuses the placenta, the cells in the endovascular compartment would be relatively similar to that of PB compared to cells within the TIS compartment. In contrast, we found that cell profiles within the EV compartment were dramatically different from those in PB and both were different from the TIS compartment. The abundance of B cells and CD4+ and CD8+ T cells differed between compartments but these cells co-localized to the same positions in the UMAPs, suggesting largely homogeneous populations of adaptive immune cells across compartments. In contrast, neutrophils, mononuclear phagocytes, basophils, and NK cells varied in their UMAP position, suggesting compartment-dependent differences in innate immune cell states (Figure 1E). For example, a NK-cell sub-cluster that is prominent in PB is absent in the placenta, and basophils were only observed in the TIS compartment. These data indicate that placental microenvironments yield both inter- and intra-organ differences in immune profiles.

To quantify compartment-specific proportions of each cell type, we calculated cell frequency as a fraction of total immune cells within each compartment (Figure 1F). Mononuclear phagocytes were the predominant cell type in both TIS and EV. Neutrophils were enriched in TIS and B cells were slightly enriched in EV. PB contained more adaptive immune cells since it contained the highest proportions of T and B cells. These intercompartmental differences were further evaluated by linear discriminant analysis (Figures 1G and S1I), where the clusters were split by organ and by compartment. Surprisingly, EV was quantitatively more similar to TIS and strikingly dissimilar to PB. To confirm these results, we employed a common ecology-based approach to quantify dissimilarity between environments (Bray-Curtis Dissimilarity). Species identifiers, typically used in these beta diversity calculations, were replaced with cell type identifiers (Horowitz et al., 2013). Beta diversity of compartmental immune composition revealed a higher similarity between EV and TIS than between EV and PB (Figure 1H).

Together these data confirm robust cell identification and the ability to quantitatively compare maternal peripheral blood and two placental microenvironments during pregnancy. The results reveal the extent to which the immune profiles of the maternal blood in the periphery and that which is filtered and perfused into the placenta diverge, and highlight the novel finding that the EV space within the placenta is a unique immune-cell niche, differentiated from both the TIS and cells circulating in the maternal periphery.

### Temporal emergence of fetal immune cells at the maternal-fetal interface

In humans and mice, placentation precedes fetal immune-cell development. For example, mature α/β T cells emerge in humans at the end of the first trimester and only emerge in the final days of gestation in the mouse with the primary expansion of this subset occurring postnatally (Mold and McCune, 2012). Implantation and early placentation occur much earlier in gestation and the placenta continues to mature until birth (Li et al., 2020; McGrath et al., 2015; Msallam et al., 2020). Due to this timeline, fetal immune-cell characterization has largely been limited to the fetal body and, other than fetal macrophages, only maternal immune cells have been examined in the placenta. Both maternal and fetal immune cells are found in umbilical cord blood (Jeanty et al., 2014), among other tissues due to microchimerism (Kinder et al., 2017), and it would be reasonable to expect the immune cells within the placenta to contain increasingly more fetal immune cells as gestation progresses.

In our gestational immune monitoring platform (Figure 1), maternal cells only express CD45.2 while fetal cells express both CD45.1 and CD45.2. This distinction was used to re-cluster fetal cells using UMAP dimension reduction with Leiden clustering on data pooled for all gestational times. The combined data from E10.5, 12.5, 14.5, and 18.5 placentas defined distinct clusters for fetal mononuclear phagocytes, neutrophils, eosinophils, B cells, T cells, and a CD45.1-positive cluster we were unable to classify (Figures 2A and S2A). We metaclustered and classified these cell types based on their expression of common lineage markers (Figures 2B and S2B). Major fetal immune-cell types were also confirmed using a manual gating strategy (Figure S2D). We found that fetal neutrophils and mononuclear phagocytes spanned multiple regions in the UMAP, suggesting significant heterogeneity in those fetal populations. All neutrophils appeared to exhibit similar Ly-6C, Ly-6G, and CD11b expression levels, but MHC-II was only expressed by cells within a distinct sub-cluster (Figures 2B and S2B). Mononuclear phagocytes showed similar CD68 expression for all cells, but Ly-6C and CD11b were differentially expressed and only one sub-cluster expressed F4/80 (Figures 2B and S2B). Mononuclear phagocytes represented more than 50% of the fetal immune cells present in the placenta in the pooled data (Figure 2C, red heatmap) but these immune cells made up only ∼0.1%–0.8% of all immune cells (Figure 2D).

**Figure 2.**
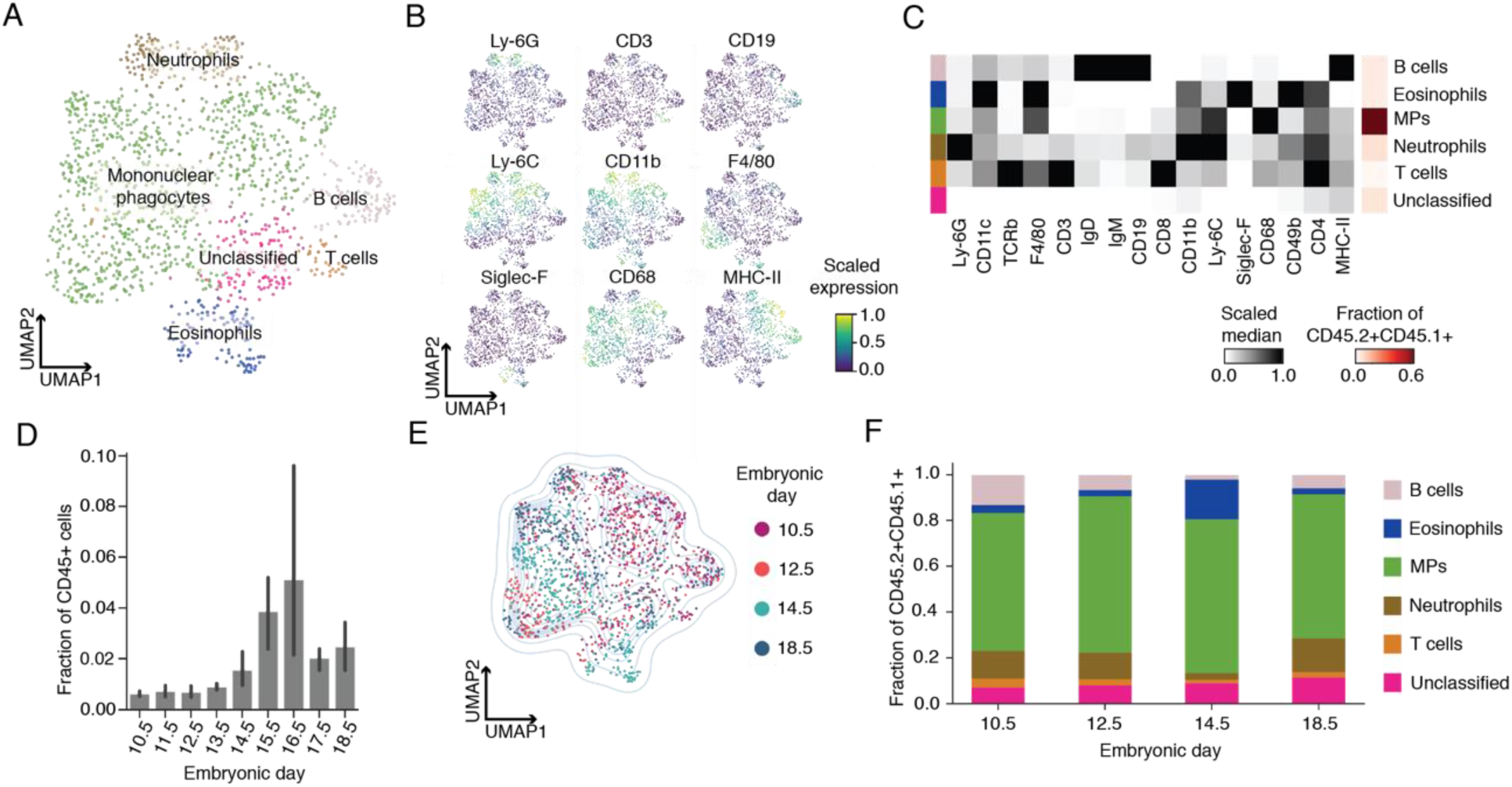
Fetal immune-cell characterization at the maternal-fetal interface. (A) Composite UMAP graph of fetal immune (CD45.2+CD45.1+) cells in placenta across E10.5 (n = 3), 12.5 (n = 3), 14.5 (n = 3), and 18.5 (n =3). (B) Scaled cellular median intensity of immune lineage markers. (C) Scaled median expression of protein markers used for Leiden clustering across fetal immune-cell types identified. Last column represents cell fraction relative to total fetal immune cells. (D) Fraction of fetal immune cells relative to all immune cells in decidua and placenta across E10.5 to 18.5. The number of samples per embryonic day used for this panel are as follows: 10.5 (n = 3), 11.5 (n = 4), 12.5 (n = 10), 13.5 (n = 3), 14.5 (n = 7), 15.5 (n= 3), 16.5 (n = 3), 17.5 (n = 3), and 18.5 (n = 5). (E) Composite UMAP graph of fetal immune cells colored by embryonic day. (F) Fraction of fetal immune cells at E10.5, 12.5, 14.5, and 18.5.

When data was evaluated for temporal changes, we noted a substantial increase in fetal cells over time (Figure 2D). To more clearly define the temporal changes in fetal immune composition at the maternal-fetal interface, the immune cells in the pooled data UMAP were colored by gestational day (Figure 2E). We found that the upper right quadrant of the UMAP (containing a subset of neutrophils and mononuclear phagocytes) was enriched with cells from E10.5 and 12.5, while the remaining quadrants featured more cells from E14.5 and 18.5. Batch differences are not a concern here because these samples were processed and analyzed simultaneously (see Methods). Interestingly, the MHC-II-expressing cells (Figures 2B and S2B), which overlap with the entire B-cell population and a single neutrophil and mononuclear phagocyte sub-region, come from E10.5 and 12.5. The F4/80-expressing mononuclear phagocytes also appeared to overlap with cells from E10.5 and 12.5. Surprisingly, we observed a stable immune profile across embryonic days 10.5, 12.5, 14.5, and 18.5 with the exception of a notable expansion of fetal mononuclear phagocytes at E14.5 (Figure 2F). When neutrophil and mononuclear phagocytes were segregated into their respective subpopulations (Figures S2A and S2B), we observed a dynamic fetal immune profile that shows the emergence of distinct neutrophil and mononuclear phagocytes over time (Figure S2C). These data suggest fetal immune cell profiles, proportions of each cell type, remain relatively stable, perhaps to maintain homeostasis, but cells also exhibit temporal transitions in phenotype within the myeloid lineages, potentially representing maturation or unique cell states that fulfill distinct functions at different times in gestation. Taken together, the relative abundance of fetal mononuclear phagocytes suggests a significant role in the placenta, especially later in gestation.

### Innate immune cell flux parallels maternal-fetal interface molecular dynamics

Pregnancy requires well-controlled tolerance mechanisms that can be perturbed in many ways by disrupting the balance of maternal immune-cell profiles. Mice with no adaptive immune system still get pregnant and have normal numbers of healthy offspring (Burke et al., 2011; Guleria et al., 2005), but the depletion of NK cells, dendritic cells, macrophages, neutrophils, or mast cells can result in failed implantation, placentation, or progression of a healthy pregnancy (Schumacher et al., 2018). In fact, a number of tolerogenic mechanisms have been proposed regarding the innate immune system’s contributions at the maternal-fetal interface (Ander et al., 2019; Arck and Hecher, 2013; Chabtini et al., 2013; Mizugishi et al., 2015; PrabhuDas et al., 2015; Schumacher et al., 2018; Trowsdale and Betz, 2006).

The mature mouse placenta is established at E10.5 and continues to grow in size and complexity until E18.5 (Figure 3A). To document if and how immune cell profiles change over time, we first asked how the immune profile changes across placental development at the level of the transcriptome. We leveraged publicly available mouse microarray gene expression data of 44 dissected fetal placenta and maternal decidua samples that spanned the lifetime of the mouse placenta (Knox and Baker, 2008). Our analysis revealed 436 immune-related and 190 vascular-related genes that significantly changed before and after E14.5 (Figure 3B, Tables S2 and S3). This molecular switch observed in late mid-gestation among genes related to vasculature coincides with the establishment and expansion of maternal-fetal exchange in the placenta. Genes related to immune-cell type and regulation also exhibited a switch at this time.

**Figure 3.**
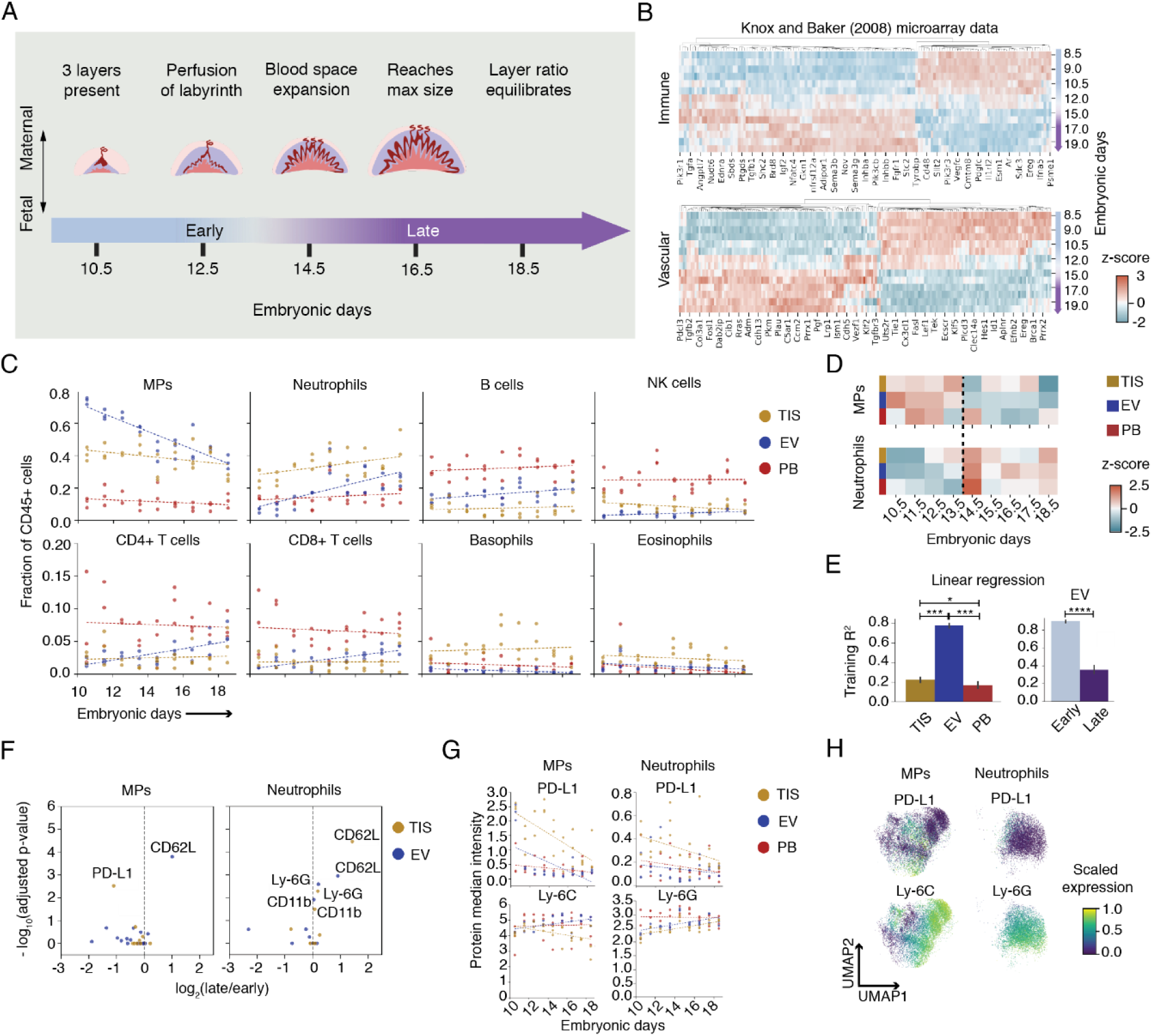
Mononuclear phagocytes and neutrophils define the gestational immune dynamics at the maternal-fetal interface. (A) Stages of placental development throughout the last half of mouse gestation. (B) Publicly available placental microarray data from Knox and Baker (2008) analyzed for immune- and vascular-associated genes that significantly changed between embryonic days 8.5 and 15.0. (C) GEE was used to fit a linear model to maternal immune cell fractions comparing TIS, EV, and PB from E10.5 to 18.5. (D) Cell fraction of maternal MPs and neutrophils across embryonic days tested, colored by z-score. All days shown have n = 3, except for E12.5 which has n = 2. (E) Training R^2^ of linear regression across each compartment based on cell fractions across embryonic days (n = 26 per compartment). Cell fractions of EV compartment were split into early (E10.5–13.5, n = 11) and late (E14.5–18.5, n = 15) and linear regression was run independently for the early and late periods. *p ≤ 0.05, ***p ≤ 0.001, ****p ≤ 0.0001 (one-way ANOVA for comparing compartments, unpaired t test for early and late stages). (F) Volcano plots of protein median intensity in MPs and neutrophils between early and late periods across TIS and EV. Only proteins with significant adjusted p-values are shown. (G) Transformed median intensity of PD-L1 and Ly-6C in MPs, and PD-L1 and Ly-6G in neutrophils were fitted with GEE to compare compartments across embryonic days. (H) Composite UMAP plots of MPs and neutrophils colored by scaled expression of protein markers.

To determine if the switch in transcriptional signatures reflects changes in immune cell profiles, mass cytometry data was re-examined without pooling timepoints and cell-type frequency was calculated for PB, EV, and TIS compartments as a fraction of total immune cells. We performed linear regression to determine if gestational day and immune composition in each compartment are correlated (Figure 3C, Table S4). Our data showed a significant linear effect between gestational day and fraction of immune cells for mononuclear phagocytes (p < 0.0001), neutrophils (p < 0.0001), CD4+ T cells (p = 0.049), and CD8+ T cells (p = 0.029) in the EV compartment, and only for neutrophils in PB (p = 0.0298). Interestingly, in addition to the significant and pronounced linear relationship between day and EV immune profile for both mononuclear phagocytes and neutrophils, these two cell types also exhibited opposite trends. Mononuclear phagocytes showed a decreasing trend while neutrophils showed an increasing trend in the EV (Figure 3C). The immune cells in the TIS compartment appear to exhibit nonlinear trends with respect to gestational day, and thus no significant linear effect could be found.

Interestingly, even though mononuclear phagocytes are temporally dynamic, they are consistently enriched in the EV compartment throughout gestation (Figure S3A). In contrast, neutrophils were TIS-biased at early timepoints and then showed no bias at later timepoints (Figure S3A). Mononuclear phagocytes and neutrophils remained the most abundant cell types in the placenta across gestation (Figure 3C), suggesting that these two innate immune-cell populations could play important roles in innate immune homeostasis in the placenta throughout gestation.

To highlight differences in immune-cell contribution over time, we calculated the z-score of cell-type abundance across embryonic days within each compartment (Figure 3D).

Interestingly, both neutrophils and mononuclear phagocytes exhibit a mutually exclusive switch between E13.5 and 14.5 in EV, suggesting a coordinated and reciprocal change in relative abundance. Linear regression analysis was then used to determine if embryonic day could be predicted by immune-cell abundance (Figures S3B and S3C). While PB and TIS performed poorly, EV showed reliable gestational dynamics that predicted gestational age (Figure S3F). Neutrophil and mononuclear phagocyte abundance alone was also able to accurately predict gestational day (Figures 3E, S3G, and S3H). To determine whether this predictability was consistent across gestation, we analyzed early and late gestation independently for all immune cells (Figures S3D and S3E) and only neutrophils and mononuclear phagocytes (Figures S3I and S3J). E14.5 marks a rapid expansion of maternal blood space and maternal-fetal exchange to support rapid fetal growth until parturition (Coan et al., 2004), prompting us to define E10.5-13.5 as early and E14.5-18.5 as late. Using all immune cells, we found that early and late gestational timepoints mapped equally well (Figure S3F, EV). Interestingly, when only using neutrophil and mononuclear phagocyte abundance, we found that early gestational timepoints were mapped 2.5- fold (p < 0.0001) more reliably than late gestational timepoints (Figure 3E, EV). This suggests that the relationship between these two cell types is critical in early stages of pregnancy.

In summary, we show pronounced molecular and cellular transitions that occur between E12.5 and E14.5, which correspond to the timing of rapid blood space expansion in the placenta. The endovascular mononuclear phagocyte and neutrophil abundance shows strong correlation with gestational dynamics and tissue remodeling at the maternal-fetal interface. These correlations are most robust in early gestation. Furthermore, our data suggest that the more pronounced changes in EV cell profiles might be a significant contributor to the transcriptomic changes for immune genes in the placenta during the molecular switch observed in the Knox and Baker dataset (Knox and Baker, 2008)(Figure 3B).

### Mononuclear phagocytes and neutrophils distinguish unique states in early and late **gestation**

Placental architecture changes throughout gestation, first to build a placenta and then to keep up with fetal metabolic demands. The placenta microenvironment is also likely to undergo dynamic changes that may influence mononuclear phagocyte and neutrophil functional states. Neutrophils and mononuclear phagocytes were examined in early and late gestation in PB, EV, and TIS compartments. We identified two molecules in mononuclear phagocytes and three molecules in neutrophils that were differentially expressed between early and late gestation (Figure 3F). CD62L, important for the capture and rolling of leukocytes on activated endothelium, increased expression on endovascular mononuclear phagocytes and neutrophils and tissue-localized neutrophils, suggesting an increase in mobility and recruitment in late gestation. Interestingly, immune checkpoint protein PD-L1 on tissue-localized mononuclear phagocytes significantly decreased over time, suggesting less need for immune suppression in late gestation, as parturition nears.

Given the increasing interest in myeloid expression of checkpoint proteins (Strauss et al., 2020), we applied linear regression to further evaluate lineage markers and PD-L1 expression across gestation on both mononuclear phagocytes and neutrophils (Figure 3G, Table S5). Ly-6C is a marker frequently used to distinguish functional subsets of myeloid lineage cells with high expression indicating recent egression from bone marrow, a proinflammatory program, or the ability to remain as a monocyte without differentiating into macrophages in tissue (Guilliams et al., 2018). Our data showed no significant linear effect between gestational day and Ly-6C expression in mononuclear phagocytes for any compartment, but the effect in TIS almost reached significance (p = 0.055). In contrast, gestational day did have a significant effect on the decreasing trend observed in PD-L1 expression over time in EV (p = 0.022) and TIS (p < 0.0001) compartments, which suggests a response to changing needs between early and late gestation. We also applied the same model to calculate the average median expression of Ly-6C and PD-L1 on mononuclear phagocytes throughout gestation for each compartment. While Ly- 6C expression did not significantly differ between PB and EV, when PB and TIS were compared, the average median Ly-6C expression was 1.11-fold lower in TIS (p = 0.004; Figure 3G). In contrast, PD-L1 expression was 0.25-fold higher in TIS compared to PB (p < 0.0001; Figure 3G). The lower Ly-6C expression and higher PD-L1 expression among mononuclear phagocytes in TIS compared to PB suggest phenotypic adaptation to the placental microenvironment.

We repeated these analyses for neutrophils and found that the effect of gestational day on the increase in Ly-6G expression over time was significant for EV (p = 0.002) and TIS (0.003), but not PB (Figure 3G). Increased Ly-6G expression, a marker of maturity (Xie et al., 2020), suggests neutrophil maturation or more mature neutrophils in the placenta during late gestation. There appeared to be a downward trend in PD-L1 expression on neutrophils across all compartments, but no significant linear effects were found. We next calculated the average median expression of Ly-6G and PD-L1 on neutrophils throughout gestation for each compartment. Ly-6G expression was 1.09-fold lower in EV (p = 0.0001) and 1.16-fold lower in TIS (p < 0.0001) compared to PB. PD-L1 expression was 0.52-fold higher in TIS compared to PB (p < 0.0001). Tissue-localized neutrophils exhibited significantly lower Ly-6G expression and higher PD-L1 expression than peripheral blood neutrophils. These data suggest an inverse relationship between neutrophil maturity and immunosuppressive potential, which aligns with observations made in the tumor microenvironment (Mackey et al., 2019). In both cell types, maturation marker expression tended to be lowest in tissue-localized cells, while PD-L1 expression tended to be highest among tissue-localized cells, suggesting specialization and compartment-specific phenotypes.

Given the significant effects gestational day had on protein expression, either global changes or a temporal flux of phenotypic subsets could be contributing factors. For example, the overall decreasing trends of tissue PD-L1 in both cell types could suggest the contraction of a PD-L1 subset in late gestation or a global decrease in PD-L1 expression. To decipher this, we examined the expression patterns of Ly-6C and PD-L1 on a mononuclear phagocyte-restricted UMAP, and Ly-6G and PD-L1 on a neutrophil-restricted UMAP (Figure 3H). The mononuclear phagocyte UMAP showed a substantial range in Ly-6C expression and multiple PD-L1-positive regions. The neutrophil UMAP showed a gradient of increasing Ly-6G expression and a single PD-L1-positive region. The differential expression of this limited number of proteins expands the possibility of a diverse myeloid compartment that remodels as a function of gestational day. Thus, these data demonstrate that mononuclear phagocytes and neutrophils change in their cell states between early and late gestation via regulatory proteins, forming compartment-specific, temporally-restricted subsets.

### Diverse and dynamic PD-L1+ mononuclear phagocytes are unique to the maternal-fetal interface

Monocytes have traditionally been dismissed as undifferentiated precursors to dendritic cells and macrophages, resulting in their diversity and function being under-explored in pregnancy. To assess mononuclear phagocyte (MP) functional heterogeneity at the maternal-fetal interface, the MPs identified in Figure 1B were re-clustered using only IdU (active S-phase), CD11c, F4/80, CD86, CD80, CD64, CD68, and MHC-II markers (Figures 4A and S4A). We identified 11 subsets of MPs and assigned functional or phenotypic identities to each subset based on their marker expression (Figures S4B and S4C). Eight out of 11 subsets were present in all three compartments (Figures S4D and S4E). We metaclustered subsets due to phenotypic similarity, resulting in 8 distinct subsets-seven of which were present in all three compartments (Figure 4A). Subsets were confirmed using canonical gating strategies where additional cell subsets could be resolved by CD11c expression (Figure S4F).

**Figure 4.**
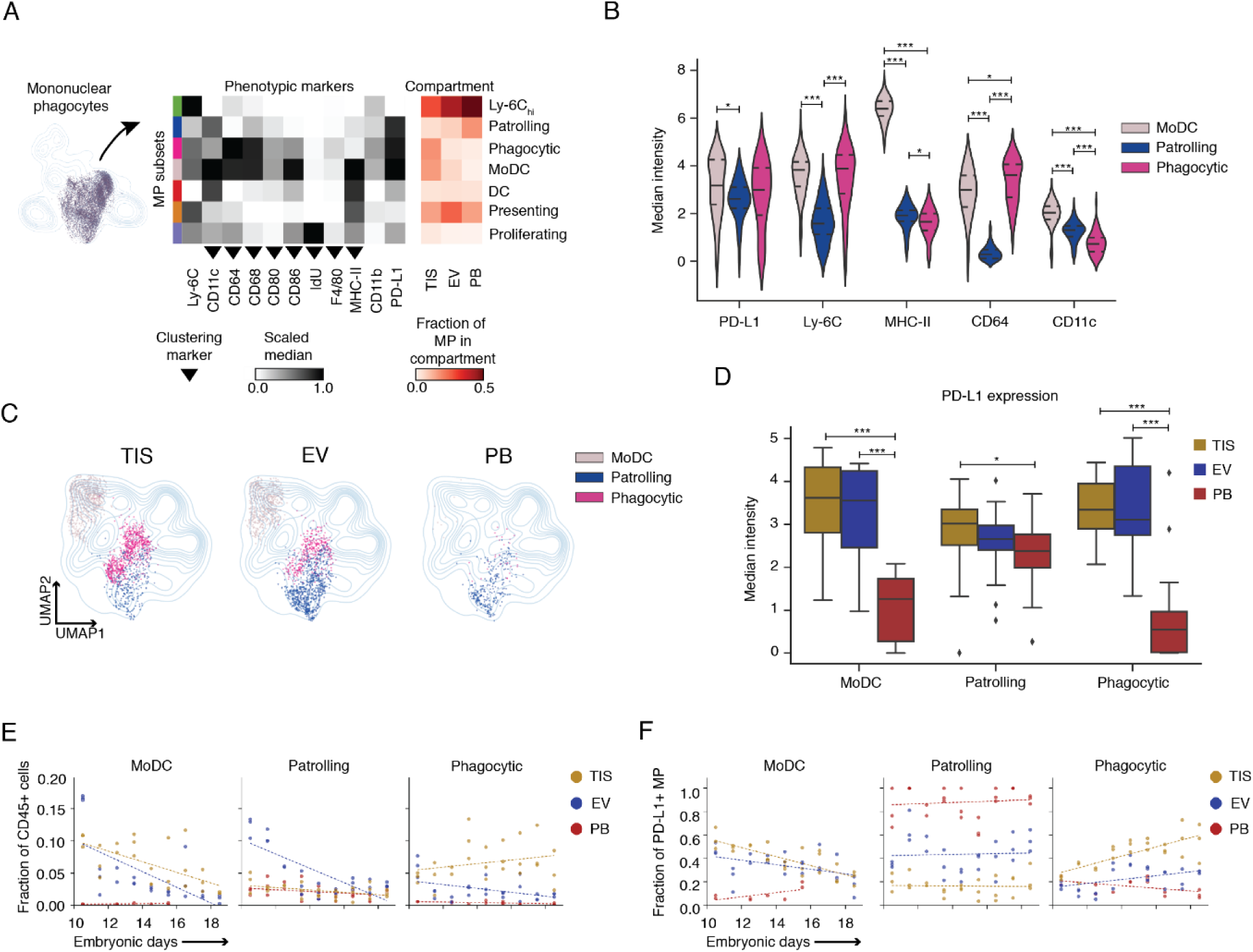
Phenotype specialization and temporal regulation of mononuclear phagocytes to placenta microenvironment. (A) Mononuclear phagocytes were reanalyzed by UMAP and Leiden clustering. Scaled medians of marker expression show seven distinct MP subsets. The last three columns of the heatmap show the fraction of each MP subset relative to all MPs in a given compartment (n = 26 per compartment). (B) PD-L1+ MP subsets were analyzed by their distinguishing phenotypic markers shown as arcsinh transformed median intensities. (C) UMAP plots showing distribution of the three PD-L1+ MP subsets across compartments. (D) Arcsinh transformed PD-L1 median intensity of moDC, patrolling, and phagocytic subsets across three compartments. (E) GEE was used to fit a linear model to fractions of moDC, patrolling, and phagocytic MP subsets relative to maternal immune cells comparing TIS, EV, and PB from E10.5 to 18.5. (F) GEE was used to fit a linear model to cell fractions of PD-L1+ MP subsets out of all PDL1+ MPs comparing TIS, EV, and PB from E10.5 to 18.5. For (B) and (D), significance is shown as *p ≤ 0.05, ***p ≤ 0.001 (one-way ANOVA per marker in (B) and per cell type in (D)).

MPs had a high expressing Ly-6C subset occupying up to ∼50% of the mononuclear phagocyte population across compartments, suggesting a majority monocyte population (Figure 4A, red heatmap). We could further classify mononuclear phagocytes by thresholding Ly-6C expression and dividing the population into Ly-6C (hi)gh, (int)ermediate, and low which are traditionally used to categorize monocytes as classical, intermediate, and non-classical monocytes, respectively (Figure S4I). Subsets that would traditionally be classified as classical or non-classical included notable fractions of all three (Figure S4J). This suggests that the conventional definition of monocytes is splitting some populations, while conflating others, potentially obscuring monocyte biology at the maternal-fetal interface. Since this population is not restricted to monocytes, it is important to note that the Ly-6C low-cells may include both non-classical monocytes and non-monocytes.

Surprisingly, the immune checkpoint ligand, PD-L1, was highly expressed in three subsets (Figure 4A). We evaluated whether these PD-L1 subsets were distinct by examining their co-expression of Ly-6C, MHC-II, CD11c, and CD64 (Figure 4B). The highest CD64-expressing, a molecule for phagocytosis, PD-L1 subset was named “phagocytic” and the highest MHC-II-expressing subset was named “moDC”, as it is most likely a monocyte-derived dendritic cell. The remaining PD-L1 subset was named “patrolling” as it was low in Ly-6C, MHC-II, CD64, and adhesion protein CD11c, indicating surveillance. These differential expression profiles within the PD-L1-expressing MPs suggest subfunction diversity. Interestingly, frequency data showed phagocytic and moDC also had a TIS enrichment (Figure 4A, red heatmap). We next interrogated their compartmental distribution. UMAP contour maps of merged data were overlaid with each compartment’s PD-L1 subsets. We found patrolling mononuclear phagocytes were present across all compartments, while moDC and phagocytic MPs, like most subsets, were largely restricted to the placenta itself (Figures 4C and S4D).

Intravascular Ly-6C low monocytes have been shown to mediate the disposal of necrotic endothelial cells during inflammation (Auffray et al., 2007; Carlin et al., 2013), whereas extravascular Ly-6C low monocytes promote angiogenesis and matrix remodeling (Nahrendorf et al., 2007). Immune cell plasticity allows them to adapt to tissues and microenvironments. Since patrolling mononuclear phagocytes were present in all three compartments, we evaluated how they changed phenotypically as they moved from PB to the placenta. The patrolling mononuclear phagocytes in TIS had significantly higher PD-L1 median expression than those in PB (1.3-fold, p = 0.037; Figure 4D), consistent with patterns observed in Figure 3G. This trend was even more pronounced in moDC (3.2-fold (TIS to PB), p = 0.001) and phagocytic (3.7-fold (TIS to PB), p = 0.001), although cells in these subsets are less abundant in PB. This upregulation in PD-L1 expression as the cells move from the PB to placenta further indicates that MPs acquire microenvironment-dependent specialization.

We next asked if these specialized subsets emerged at different points of gestation. MoDC and phagocytic MPs were enriched in the placenta and had a significant TIS bias compared to PB (moDC: p < 0.0001, phagocytic: p < 0.0001) for most of gestation, whereas patrolling MPs were only placenta enriched at E10.5 and E12.5 after which their frequency in EV became comparable to PB (Figure 4E, Table S6). MoDC and patrolling MPs exhibited a short-lived EV bias at E10.5, confirming previous reports (Kruse et al., 1999). MoDCs appeared to decrease over time in both TIS and EV, while phagocytic MPs appeared to increase in TIS. Linear regression analysis showed a significant linear effect between gestational day and fraction of immune cells for moDCs in EV (p = 0.0001) and TIS (p < 0.0001), patrolling MPs in PB (p = 0.035) and EV (p < 0.0001), and phagocytic MPs only in EV (p = 0.028) (Figure 4E). As a fraction of PD-L1-expressing mononuclear phagocytes, moDC consistently showed placental bias on average compared to PB (TIS: p < 0.0001, EV: p < 0.0001; Figure 4F, Table S7). Phagocytic MPs also showed a placental bias on average compared to PB (TIS: p < 0.0001, EV: p = 0.002), and the TIS bias increased over time (Figure 4F). MoDCs showed an overall decreasing trend in TIS and EV, while phagocytic MPs inversely increased. There was also a significant linear effect between gestational day and fraction of PD-L1-expressing MPs for moDCs (PB: p = 0.01, EV: p = 0.0008, TIS p < 0.0001) and phagocytic MPs (PB: p = 0.004, EV: p < 0.0001, TIS p < 0.0001) across all compartments. Patrolling MPs occupied the majority of peripheral blood’s PD-L1-expressing mononuclear phagocyte population throughout gestation and showed no significant linear effect by gestational day (Figure 4F). These data suggest that moDC and patrolling MPs might serve important regulatory roles establishing the mature placenta, while phagocytic MPs continue to play an important regulatory role throughout gestation as it potentially assists in tissue remodeling and other homeostatic processes.

In addition to the PD-L1-expressing MPs, we also tracked other subsets over time. As a fraction of all immune cells, the monocytic Ly-6Chi subset remained the most abundant MP population throughout gestation and showed a consistent EV bias (p < 0.0001) on average compared to PB (Figure S4G, Table S6). Presenting MPs show an EV bias (p < 0.0001) on average compared to PB that is pronounced between E10.5 and E15.5 with a clear decreasing trend over time. There was a significant linear effect between gestational day and presenting MPs in both EV (p < 0.0001) and PB (p < 0.0001), but not for Ly-6Chi. When we look at these subsets as a fraction of MPs, it becomes clear that the MP population becomes enriched with Ly- 6Chi MPs as gestation progresses (Figure S4H, Table S8). An enrichment of high Ly-6C- expressing monocytes could be due to more recent bone marrow emigrants being released into the periphery, or a systemic shift to proinflammatory subsets in preparation for parturition. Ly- 6Chi is the most abundant subset in PB within the MP population, and while clearly high and increasing over time in all compartments, exhibits a PB bias compared to the placenta on average (EV: p = 0.02, TIS: p < 0.0001) and significant linear effect in PB (p = 0.02; Figure S4H). The presenting subset remains highest in the EV compartment, but the bias and decreasing trend are less pronounced. Regardless, presenting MPs showed a significant linear effect by gestational day for all compartments (PB: p < 0.0001, EV: p = 0.025, TIS: p = p < 0.0001; Figure S4H).

Since PD-L1 expression has largely been restricted to non-classical and intermediate monocytes and is used as a differentiating marker (Bianchini et al., 2019)), we also examined PD-L1 expression on the classical, intermediate, and non-classical monocytes we identified. Classical monocytes in PB indeed lacked PD-L1, but upon entering TIS, our single-cell data showed that PD-L1 expression was upregulated (E12.5: 3.5-fold, p ≤ 0.001, E14.5: 3.9-fold, p ≤ 0.001; Figure S4K). In addition to an increase in PD-L1 median intensity, there was a higher fraction of classical (high Ly-6C-expressing cells) monocytes that expressed PD-L1 in TIS compared to PB (E12.5: 4.8-fold, p ≤ 0.001, E14.5: 6.9-fold, p ≤ 0.001; Figure S4L). These data highlight that classical monocyte expression of PD-L1 is dependent on environment, and PD-L1 expression should not be used as a classical monocytes identifier in tissue microenvironments.

Collectively, these data show both increased heterogeneity of mononuclear phagocyte cell states and PD-L1 phenotypic expansions within the placenta, likely due to its unique local microenvironments compared to PB. These specialized subsets exhibit organ and compartment bias as well as unique temporal dynamics which align with structural changes at the maternal-fetal interface that necessitate regulatory mononuclear phagocytes that can adapt to and assist with extensive tissue remodeling and wound healing.

### Noncanonical neutrophils in early gestation placenta bare PD-L1 and proliferate *in situ*

While remarkably diverse in form and function, neutrophils are often characterized as an injurious monolith, commonly associated with pregnancy complications (Giaglis et al., 2016; Tong and Abrahams, 2020). Our data show neutrophils as being an abundant cell type at the healthy maternal-fetal interface throughout gestation (Figure 3C), suggesting an important homeostatic role during pregnancy. Additionally, initial protein analysis identified a subset of neutrophils expressing checkpoint protein PD-L1 (Figures 3F-3H), suggesting regulatory potential and heterogeneity. Surprisingly, the neutrophils here expressed many of the regulatory immune receptors more typically associated with MP function. Given this observation, we employed a similar approach to the MP analysis (Figure 4) to dissect neutrophil diversity by re-clustering neutrophils based on CD80, CD62L, CD40, MHC-II, PD-L1, and IdU markers (Figures 5A and S5A-S5C). Eleven subsets were generated by Leiden and found across compartments (Figures S5D and S5E), but further reduced based on phenotypic similarity.

**Figure 5.**
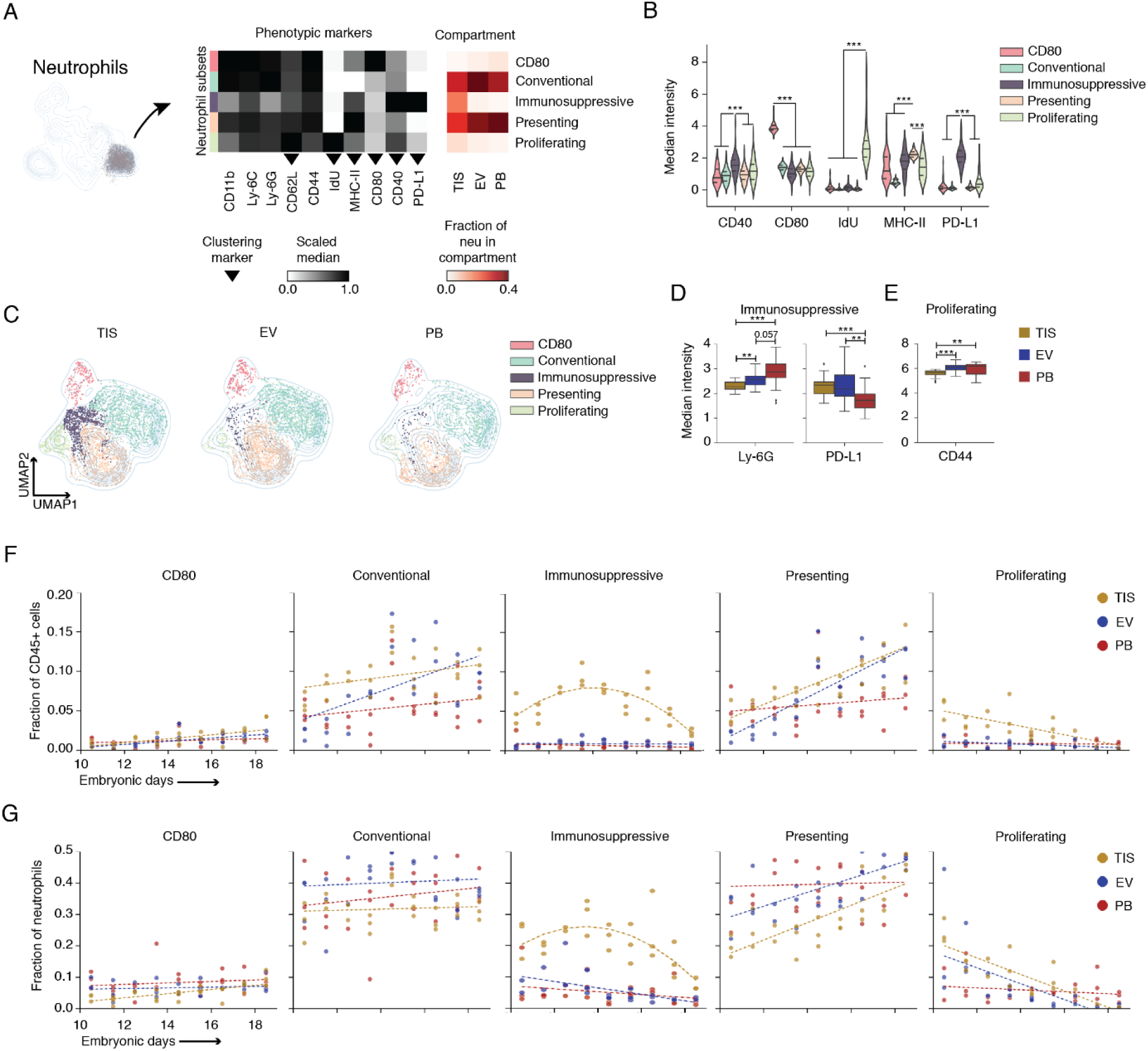
Placenta enriches for noncanonical neutrophil subsets in tissue compartment-specific manner. (A) Neutrophils were reanalyzed by UMAP and Leiden clustering. Scaled medians of marker expression shows five distinct neutrophil subsets. The last three columns of the heatmap show the fraction of each subset relative to all neutrophils in a given compartment (n = 26 per compartment). (B) Neutrophil subsets were analyzed by their distinguishing phenotypic markers shown as arcsinh transformed median intensities. (C) UMAP plots showing distribution of the five neutrophil subsets across compartments. (D) Arcsinh transformed Ly-6G and PD-L1 median intensity of the immunosuppressive subset across three compartments. (E) Arcsinh transformed CD44 median intensity of the proliferating subset across three compartments. (F) GEE was used to fit a linear model to fractions of CD80, conventional, presenting, and proliferating subsets relative to maternal immune cells comparing TIS, EV, and PB from E10.5 to 18.5. GEE was applied to fit a quadratic model to the immunosuppressive subset. (G) GEE was used to fit a linear model to CD80, conventional, presenting, and proliferating subsets relative to total neutrophils comparing TIS, EV, and PB from E10.5 to 18.5. GEE was applied to fit a quadratic model to the immunosuppressive subset. For (B), (D), and (E), significance is shown as **p ≤ 0.01, ***p ≤ 0.001 (one-way ANOVA per marker in (B) and per cell type in (D) and (E)).

Five distinct subsets were identified and classified as “presenting”, “immunosuppressive”, “proliferating”, “CD80”, and “conventional” neutrophils due to their respective MHC-II, PD-L1, IdU, CD80, and canonical expression profiles (Figure 5B). Further, these subsets were confirmed and easily identified using canonical gating strategies (Figure S5F). We found subsets of relatively high MHC-II-expressing cells within the immunosuppressive, proliferating, and CD80 subsets (Figures S5B and S5C). We also found a subset of CD40-expressing cells in both the conventional and presenting subsets (Figures S5B and S5C). CD40 is a member of the tumor necrosis factor receptor family and is important in co-stimulation, anti-tumor immunity, and immune regulation upon binding to its ligand, CD40L (Foey et al., 2001; Portillo et al., 2012; Rakhmilevich et al., 2012). PB was enriched with the CD40-negative fraction of the conventional and presenting subsets, whereas the placenta had uniform distribution of the entire conventional and presenting subsets (Figures 5C, S5B, and S5D). While notable, none of these observations warranted further classification as distinct subsets (see Methods).

Merged data from all embryonic days showed conventional and presenting subsets occupied the majority of the neutrophil population (Figure 5A, red heatmap). Furthermore, potentially immunosuppressive and proliferating neutrophils appeared to preferentially populate TIS (Figures 5A, red heatmap, and 5C). The remaining subsets, while present in all compartments, were more enriched in EV and PB. Since immunosuppressive and proliferating subsets demonstrated a TIS bias, but were not TIS restricted, we asked how these two subsets differed phenotypically in EV and PB. As with the MPs (Figure 4), as the immunosuppressive (PD-L1) neutrophils moved from PB to EV, their PD-L1 expression increased 1.4-fold (p = 0.006) and its Ly-6G expression decreased 1.12-fold (p = 0.06; Figure 5D). There was a further 1.1-fold (p = 0.01) decrease in Ly-6G as PD-L1-expressing neutrophils moved from EV to TIS. Interestingly, immune cells in the tumor microenvironment can regress in maturity as a form of immune regulation (Mackey et al., 2019) and Ly-6G has been used to assess the level of maturity for neutrophils (Xie et al., 2020).

Mature neutrophils are mitotically inactive with cell-cycle arrest, but in cancer, neutrophil granulopoiesis can occur outside of the medullary spaces of the bone marrow (Mackey et al., 2019). With the numerous parallels between tumor immunology and tolerizing immune mechanisms in pregnancy, it is reasonable to posit that neutrophil precursors could be seeding the placenta and participating in local granulopoiesis. This interesting subset of CD40 baring, proliferating neutrophils had decreased CD44 expression (1.07-fold, p = 0.005) as it moved from PB to TIS (Figure 5E) and was significantly enriched in the placenta (Figures 5A, red heatmap, and 5C). CD44 is a protein involved in attachment to and rolling on endothelial cells as well as homing to peripheral lymphoid organs and sites of inflammation. As we observed among mononuclear phagocytes, this upregulation in functional protein expression as these subsets moved from the PB to the placenta suggests specialization for the placental microenvironment.

As with MPs (Figure 4), we next interrogated the population dynamics across gestation. Conventional and presenting neutrophils remained the most abundant across compartments throughout gestation and showed placenta bias at late timepoints (Figure 5F, Table S9). On average, the conventional neutrophil fraction was higher in the placenta compared to PB (EV: p = 0.01, TIS: p < 0.0001), as was the presenting neutrophil fraction (TIS: p = 0.001). Immunosuppressive neutrophils (p < 0.0001) and proliferating neutrophils (p < 0.0001) demonstrated TIS bias on average compared to PB throughout gestation. CD80 neutrophils were consistently the least abundant subset throughout gestation. Conventional, presenting, and CD80 neutrophils showed an increasing trend in all compartments, while proliferating neutrophils seemed to have a decreasing trend, most pronounced in TIS. To determine if any of these trends can be explained by gestational day, we performed linear regression. There was a significant linear effect between gestational day and fraction of immune cells for CD80 neutrophils in all compartments (PB: p = 0.02, EV: p = 0.0008, TIS: p = 0.006), conventional neutrophils in PB (p = 0.03) and EV (p = 0.005), presenting neutrophils in EV (p < 0.0001) and TIS (p = 0.0009, and proliferating neutrophils in TIS (p < 0.0001). Tissue-localized immunosuppressive neutrophils exhibited a negative parabolic-shaped trend during gestation and required a nonlinear fit. There was a significant quadratic effect between gestational day and immunosuppressive neutrophils in TIS (p < 0.0001), showing an initial rise between E10.5 and E13.5 and then a gradual decrease at later timepoints.

As a fraction of neutrophils, conventional and presenting neutrophils were the dominant subsets in PB and EV for most of gestation (Figure 5G). While immunosuppressive (p < 0.0001) and proliferating (p = 0.001) subsets maintained a TIS bias when compared to PB (Figure 5G, Table S10). There were only significant linear effects between gestational day and fraction of neutrophils for presenting neutrophils in EV (p = 0.009) and TIS (p = 0.001) and proliferating neutrophils in EV (p = 0.007) and TIS (p < 0.0001). The nonlinear relationship between tissue-localized immunosuppressive neutrophils and gestational day reappeared, exhibiting a significant quadratic effect (p < 0.0001). Neutrophil subsets showed clear and distinct trends during gestation, demonstrating notable temporal dynamics that suggest neutrophil responsiveness to a changing environment during pregnancy.

### Response to Poly(I:C) challenge is gestationally dependent, reducing placental PD- L1+ cells

Systemic immune activation during pregnancy, especially in early gestation, can dysregulate the mechanisms at the maternal-fetal interface that maintain tissue homeostasis and tolerance, increasing the risk of pregnancy and post-natal complications (Yockey and Iwasaki, 2018). There is a critical need to understand how activation of the maternal innate immune system, especially by viral infections, alters the immune state of the placenta. Using our gestational immune monitoring framework, we performed a systemic perturbation on pregnant dams at E12.5 or E14.5 to identify significant changes that occur across the transition from early to late gestation identified in Figure 3 and to understand how this timing might impact the immune response. Specifically, we examined the extent to which the immune profiles of all three compartments could change two hours after an intraperitoneal injection of the viral antigen mimic Poly(I:C) (Figure 6A).

**Figure 6.**
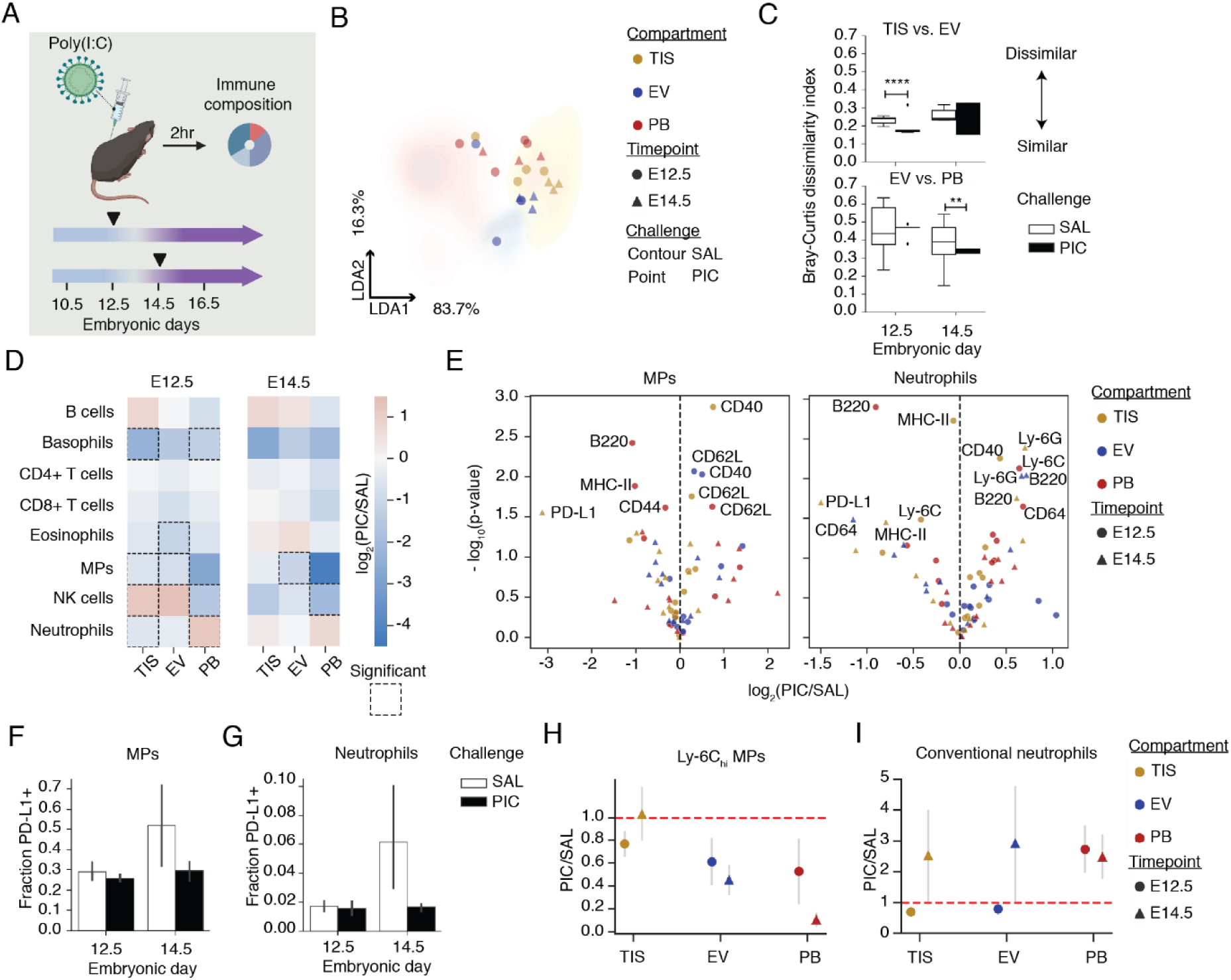
Immune response to systemic perturbation is dependent on gestational day. (A) Experimental set up of systemic maternal immune challenge with Poly(I:C) (PIC). The number of animals used for each condition are as follows: baseline E12.5 TIS (n = 10), EV (n = 10), PB (n = 6); baseline E14.5 TIS (n = 7), EV (n = 7), PB (n = 7); PIC E12.5 TIS (n = 4), EV (n = 4), PB (n = 4); PIC E14.5 TIS (n = 3), EV (n = 3), PB (n = 3). (B) LDA based on maternal immune cell fractions in each compartment. Contour plots show baseline, saline (SAL) samples, and points represent PIC samples. Circles represent E12.5 samples, and triangles represent E14.5 samples. (C) Bray-Curtis dissimilarity based on maternal immune-cell fractions in each compartment and the given immune challenge. **p ≤ 0.01, ****p ≤ 0.0001 (unpaired t test). (D) For E12.5 and 14.5, maternal immune-cell fraction was compared by taking the log2 ratio of PIC over SAL. Significant changes between challenges are encased by a dotted line (p < 0.05). P values are uncorrected for multiple comparisons. (E) Volcano plots of protein median intensity changes in MPs and neutrophils between SAL and PIC across TIS, EV, and PB. Shape of the points represent embryonic day. Only proteins with significant changes between SAL and PIC are shown. P values are uncorrected for multiple comparisons. (F) PD-L1+ fraction of MPs at E12.5 and 14.5 with or without PIC challenge. (G) PD-L1+ fraction of neutrophils at E12.5 and 14.5 with SAL or PIC challenge. (H) The PIC over SAL counts of Ly-6Chi MP subset and conventional neutrophil subset were analyzed across embryonic days (point shape) and compartments.

Intercompartmental differences following the systemic immune challenge were evaluated by linear discriminant analysis using cell abundance (Figure S6B), where baseline distributions are depicted as density contour maps and data points for individual Poly(I:C) treated animals overlaid as individual circles (E12.5) or triangles (E14.5) (Figure 6B). While Poly(I:C)-treated TIS composition remained largely unchanged, overlapping with TIS distributions in saline controls, the PB and EV profiles were substantially perturbed relative to saline and mapped more closely to the TIS cell profiles. This suggests a significant disruption in PB and EV populations following Poly(I:C) injection, but a relative stability in the profiles of cells in the TIS compartment. The calculated beta diversity between EV and TIS as well as EV and PB (Figure 6C) confirmed a significant increase in similarity between EV and TIS at the early timepoint (1.13-fold change, p < 0.0001), and between EV and PB at the late timepoint (1.2-fold change, p = 0.002). The increase in similarity between these compartments is due to the Poly(I:C) perturbation.

To determine the immune cell populations that mediated the Poly(I:C)-induced changes, we calculated cell frequency as a fraction of total immune cells (Figure S6A) and summarized the results with log-fold change of major cell types within PB, EV, and TIS compartments (Figure 6D). Poly(I:C) at E12.5 evoked changes in several cell types in all 3 compartments while Poly(I:C) at E14.5 had less impact. At E12.5, Poly(I:C) injection resulted in an enrichment of neutrophils and depletion of basophils and NK cells in PB, a depletion of eosinophils and mononuclear phagocytes in EV, and depletion of basophils, MPs and neutrophils with increase in NK cells in TIS. This suggests compartmentalized responses to Poly(I:C), and the placenta’s ability to continue to regulate cell access from perfusing maternal blood during a systemic immune perturbation. Coinciding with the molecular and immunological switch we see (Figure 3), we hypothesized that immune perturbations before E14.5 would lead to more disruption of the immune profile at the maternal-fetal interface than after E14.5. E14.5, in fact, had fewer changes following Poly(I:C) administration than E12.5, with only an NK-cell depletion in PB and mononuclear phagocyte depletion in EV. Even between practically adjacent embryonic days, we see responses to a perturbation being dependent on gestational age with a higher magnitude of responses occurring earlier in gestation (Figure 6D). Such observations could coincide with vulnerable periods of pregnancy, seen in both human (Allanson et al., 2010; Srinivas et al., 2006) and mouse (Carpentier et al., 2011a, 2013; Hsiao and Patterson, 2011; Vermillion et al., 2017) gestation.

To determine which MP and neutrophil subsets were most affected by Poly(I:C), we calculated their frequency as a fraction of total immune cells within each compartment. Within the MP population, patrolling MPs contracted in PB and EV on both E12.5 and E14.5; moDC and patrolling MPs contracted in TIS on both days, but phagocytic MPs only contracted in TIS on E14.5 (Figure S6C). Within the neutrophil population, immunosuppressive neutrophils contracted in TIS on E12.5 and E14.5, while the proliferating subset contracted in TIS and expanded in EV on E14.5 (Figure S6D). The disruption that Poly(I:C) exerted on MP and neutrophil subsets depended on gestational day and compartment.

Since mononuclear phagocytes and neutrophils are the innate immune sensors for pathogens, we examined how expression of functional proteins changed after Poly(I:C) injection. There were 7 significantly changed activation molecules in mononuclear phagocytes and neutrophils with only MHC-II, B220, CD40, and PD-L1 in common (Figure 6E). PD-L1 had the highest negative magnitude of difference of expression following immune perturbation in both cell types, but only at E14.5 and only in TIS. Mononuclear phagocytes had more significant molecular expression changes at E12.5 than at E14.5. Based on the number of molecules significantly changed within a compartment following the immune perturbation, mononuclear phagocytes showed the most activation in PB, whereas neutrophils showed it mostly in TIS.

Considering PD-L1-expressing MP and neutrophil subsets contracted following Poly(I:C) (Figure S6C and S6D) and PD-L1 was the most significantly changed molecule in both tissue-localized mononuclear phagocytes and neutrophils (Figure 6E), we quantified the fraction of PD- L1-expressing cells in both populations (Figures 6F and 6G). Poly(I:C) perturbation resulted in a 1.8-fold reduction of PD-L1-expressing mononuclear phagocytes at E14.5, which mostly impacted the proliferating subset (Figures 6F and S6E). Neutrophils had a 3.7-fold reduction at E14.5, which consequently reduced both the conventional and proliferating subsets (Figures 6G and S6F). Poly(I:C) also caused a decrease in the median expression of PD-L1 across every mononuclear phagocyte subset except phagocytic and presenting (Figure S6G), and every neutrophil subset except presenting in TIS at E14.5 (Figure S6H). However, Poly(I:C) administration resulted in increased PD-L1 expression among mononuclear phagocyte subsets, like Ly-6Chi, in PB and EV on both days, but substantially higher at E12.5. While the tissue compartment was largely protected (Figure 6B), it still showed rapid modulation of PD-L1 and PD-L1-expressing cells. Given the short time period, perhaps these PD-L1-expressing neutrophils and mononuclear phagocytes are being ‘mobilized’ out of the placental microenvironment by the Poly(I:C) perturbation.

Since there were temporal and compartmental differences in the PD-L1-expressing fractions of mononuclear phagocytes and neutrophils, we next interrogated how the most abundant and potentially inflammatory subpopulations responded to Poly(I:C). Ly-6Chi MPs contracted on both days in PB and EV, but notably more so at E14.5 (Figures 6H and S6C). Whereas only TIS at E12.5 saw a modest reduction. At the same time, there was a difference between the TIS and PB responses at E14.5, but not E12.5. The most abundant neutrophil subset, conventional, experienced an expansion on both days in PB, but only on E14.5 in the placenta (Figures 6I and S6D). There was a difference between the TIS and PB responses at E12.5, but not E14.5. Altogether, these findings show that even two hours after a systemic perturbation of the maternal immune system, the placenta can respond rapidly and uniquely depending on the compartment and gestational day.

Given these findings in the maternal immune compartment, we next examined how a systemic immune perturbation might impact the fetal immune compartment of the placenta. Fetal mononuclear phagocytes significantly contracted 1.4-fold (p = 0.004) in the placenta within two hours of the systemic Poly(I:C) perturbation at E14.5, but not at E12.5 (Figure S6I). Additionally, there was a significant 2.6-fold (p = 0.01) contraction in the unclassified population at E12.5, but not at E14.5. This fetal response is distinct from the maternal response, which showed a significant reduction in mononuclear phagocytes on both days. These data suggest that the fetal immune response in the placenta is also dependent on gestational day.

Overall, tissue-localized PD-L1-expressing neutrophils and mononuclear phagocytes were the most modulated subsets following the systemic immune perturbation. Even when the TIS compartment of the placenta remained largely unchanged (Figure 6D, E14.5), we see shifts away from PD-L1-baring cells (Figures 6F and 6G, E14.5) that may contribute to loss of tolerance in immune activation and allogeneic scenarios. The myeloid heterogeneity we have characterized here potentially impacts regulation in pregnancy at both steady-state and a perturbed-state. The phenotypic, compartment-specific, and temporal characterization of these subpopulations provides a foundation to understand and assess the functionally diverse contributions of myeloid cells at the maternal-fetal interface. This demonstrates a broad utility for our gestational immune monitoring approach and framework for the interrogation of immune-cell dynamics and phenotypic heterogeneity at, but not limited to, the maternal-fetal interface during pregnancy.

## DISCUSSION

Pregnancy model systems that examine immune tolerance are challenging with conventional knock-in or knock-out experiments due to the preponderance of embryonic lethal phenotypes or downstream consequences that complicate interpretation (Ingman and Jones, 2008; Schumacher et al., 2018). Instead of traditional loss-of-function experiments, we mapped the compartmental immune dynamics of the placenta over gestation to identify associated cell states, abundance, and localization relationships that could be participating in a regulatory capacity. We leveraged single-cell mass cytometry and generated broad immune-cell profiles throughout the second half of mouse gestation, after the placenta has formed. Our strategy (Figure 1) was able to capture both maternal- and fetal-derived immune cells and discern the compartment of origin for maternal immune cells, either circulating in maternal peripheral blood, anchored in the maternal endovascular (EV) space within the placenta, or residing within placental tissue (TIS) proper.

Initially, we anticipated that immune cell profiles would be relatively similar in both peripheral blood and placental EV compartments. Surprisingly, we found that maternal cell profiles in the placental endovascular space substantially diverged from those in the maternal periphery (Figures 1G and 1H). Maternal immune cells within the placental tissue showed even higher levels of specialization and our results uncover a novel placental adaptation of immunomodulatory mononuclear phagocytes (Figure 4) and neutrophils (Figure 5) that may substantially contribute to immune tolerance for the semi-allogenic placenta and fetus.

The gestational dynamics observed for both fetal (Figure 2) and maternal (Figure 3) innate immune-cell types within the placenta captured immune adaptation that shares surprising similarities to immune evasion and mechanisms being uncovered in tumor biology (Holtan et al., 2009b; Lala et al., 2021b). In particular, we highlighted PD-L1-expressing mononuclear phagocytes and neutrophils and their transience in the placental endovascular and tissue compartments as significant candidates in mediating immune tolerance in pregnancy.

More broadly, neutrophils and mononuclear phagocytes displayed inverse relationships over time which suggests a high degree of coordination within these relatively abundant placental immune cell classes in pregnancy (Figures 3C and 3D). This gestational switch, highlighted by an E12.5 and E14.5 transition, was reinforced by bulk molecular analysis (Figures 3B-3D) and aligns with vascular remodeling and blood space expansion (Figure 3A).

These gestational immune dynamics, and thus transition, could be predicted by neutrophils and mononuclear phagocytes, particularly those in the endovascular space (Figure 3E). Such tight coordination between neutrophils and mononuclear phagocytes appears critical at midgestation, and aligns with the period of high susceptibility to immune-related pregnancy complications (Allanson et al., 2010; Carpentier et al., 2011a, 2013; Hsiao and Patterson, 2011; Srinivas et al., 2006). Furthermore, the higher level of gestational coordination in the endovascular compartment compared to the placenta tissue indicates that endovascular immune cells contribute significantly to the transcriptional switch observed in the bulk molecular data (Knox and Baker, 2008)(Figure 3B).

In exploring the composition of these innate immune populations, we identified 7 distinct subsets within placental mononuclear phagocytes (Figure 4) that were not sufficiently described using conventional peripheral blood- or bone marrow-derived definitions. Thus, to better capture the heterogeneity of innate immune cell types observed at the maternal-fetal interface, we created a naming scheme meant to reflect the inferred cell function based on overall phenotype. Conventionally viewed as dendritic cell or macrophage progenitors, monocytes now have recognized immune functions without requisite differentiation (Guilliams et al., 2018; Olingy et al., 2019). Given that most mononuclear phagocyte subpopulations were placenta-enriched (Figure S4D), it is likely that the local tissue microenvironment instructs the cell state and recruitment of these rare cells to maintain specific functionality.

One critical function during pregnancy is the suppression of maternal immune cells from attacking the fetus. Among the MPs here, we identified 3 PD-L1-expressing subsets: moDC, patrolling, and phagocytic (Figure 4A). As deletion of PD-L1 in dendritic cells results in enhanced antitumor immunity (Oh et al., 2020) our data suggests that these PD-L1+ subsets could be modulating adaptive immunity and supporting placental growth. Furthermore, PD-L1 monocyte-derived subsets have been reported to support other functions related to placentation, including the early response to tissue damage (i.e. remodeling) and scavenging debris in vessels and tissues (Bianchini et al., 2019; Olingy et al., 2017). Given the broad role of MPs in immune regulation and tissue homeostasis and remodeling, it is not surprising that the abundance and localization of unique PD-L1 subsets in the placenta is strongly linked to gestation, which supports a significant role during pregnancy.

Exhibiting a similar significance to MPs in our study, neutrophils had remarkable diversity at the maternal-fetal interface with 5 subsets identified (Figure 5A). These data demonstrated neutrophil heterogeneity well beyond what is observed in the periphery and instead associated with complex immune tissues (Evrard et al., 2018; Grieshaber-Bouyer and Nigrovic, 2019; Xie et al., 2020) and the tumor microenvironment (Sagiv et al., 2015). Understudied in pregnancy, and typically classified based on morphology and a few immunophenotypic molecules, neutrophil diversity and possibly associated functions are not widely reported. Interestingly, while single-cell RNA sequencing has been applied to explore immune-cell heterogeneity at the maternal-fetal interface, granulocytes were seemingly omitted from the analysis (Vento-Tormo et al., 2018) possibly due to technical reasons including their sensitivity to isolation protocols, low RNA content, and high RNase levels (10X Genomics (2020)).

Still, the general role of neutrophils in pregnancy has been assessed en masse and related to their role in placental damage and preterm labor (Giaglis et al., 2016; Tong and Abrahams, 2020). However, neutrophil depletion during pregnancy in an attempt to abate such injury, had the opposite effect, instead exasperating it (Higashisaka et al., 2018). Thus, neutrophil heterogeneity, as depicted in our study, implies functional diversity and a balanced ecosystem where depleting or over-stimulation would disrupt this equilibrium and inhibit protective functions that might be present. Possibly playing a role in maintaining this balance is the IdU+ (i.e. actively cycling, S-phase) neutrophils we identified that decreased over gestation (Figures 5F and 5G). Sometimes seen in cancer, but not typically outside of bone marrow (Mackey et al., 2019), these proliferating placental neutrophils could be related to the increased frequency of immature granulocytes in peripheral blood during pregnancy (Blazkova et al., 2017). Given that neutrophil precursors can seed distant tissues, producing more cells in support of tumor growth and metastasis (Kowanetz et al., 2010; Wu et al., 2018), it seems that the CD40 and PD-L1- expressing proliferating subset could be expanding in the placenta to regulate the activity of T cells, macrophages and NK cells as previously reported (Rakhmilevich et al., 2012).

Along with the proliferating subset, we found an abundant and dynamic population of PD-L1+ neutrophils at the maternal-fetal interface, that could be complementing the role of previously reported immune-suppressive myeloid-derived suppressor cells (MDSCs) in the placenta (Ahmadi et al., 2019). Like tumors producing the neutrophil/monocyte chemoattractant CCL2 (Wu et al., 2018), we have previously shown that the healthy placenta releases significant amounts of CCL2 at steady state (Carpentier et al., 2011b), making it plausible that the placenta actively recruits these immature and immunosuppressive cells in addition to their local maintenance from the proliferative subset. While they have not been reported in pregnancy, the overall regulatory effects of PD-L1+ neutrophils has been shown in cancer and microbial infections (Bowers et al., 2014; Castell et al., 2019; Chun et al., 2015; de Kleijn et al., 2013; Langereis et al., 2017; Mcnab et al., 2011; Schulte-Schrepping et al., 2020). Furthermore, tumor-induced PD-L1 upregulation on neutrophils reportedly increases the neutrophil lifespan (Cheng et al., 2018), enabling them to exert their suppressive potential longer, thereby accentuating tumor growth and progression (He et al., 2015; Wang et al., 2017). Given that both IdU+ and PD-L1+ neutrophil subsets had a significant placental tissue bias and specific enrichment at earlier timepoints (Figures 5F and 5G), we speculate that they might be supporting placenta immune evasion, growth, and development, like their contributions in tumor growth and development.

Complementing numerous studies that describe an overall enrichment of the PD-1/PD-L1 axis in the placenta compared to the periphery (Gong et al., 2014; Guleria et al., 2005; Meggyes et al., 2019; Petroff et al., 2003; Veras et al., 2017), our unique implication of PD-L1-expressing mononuclear phagocytes and neutrophils that are tightly linked to gestational remodeling presents a new role for these cells and this pathway at the maternal-fetal interface.

Systemic activation of the immune system during pregnancy has been shown to increase the risk for pregnancy complications, and in the offspring, allergy/asthma and neurodevelopmental disorders (Atladóttir et al., 2010; Baud et al., 2008; Brown, 2011; Brown et al., 2000, 2004; Croen et al., 2005; Lee et al., 2015; Solek et al.; Sørensen et al., 2009; Zerbo et al., 2015). Furthermore, a preterm birth mouse model that uses a bacterial mimic to activate the maternal immune system results in an imbalance between adaptive and innate immune cells in the placenta (Arenas-Hernandez et al., 2015), suggesting the importance of a balanced compartment in maintaining a healthy pregnancy to term. Given the level of coordination and potentially regulatory nature we observed for innate immune cells at the maternal-fetal interface, it is reasonable to suspect that activation of these cells could disrupt the delicate maintenance of the placenta. Here, activating a toll-like receptor (TLR)-3 maternal innate response with the viral mimic Poly(I:C) showed loss of PD-L1 expression as one of the most pronounced effects (Figures 6E-6G). At the same time, we observed a reduction of Ly-6Chi MPs (Figure 6H) and an expansion of conventional neutrophils within the tissue compartment (Figure 6I), mimicking acute inflammation. The extent to which both the placental tissue and endovascular immune compartments could remodel within two hours of a systemic perturbation was surprising. This observation suggests an abrupt disturbance in the mechanisms necessary to maintain compositional balance.

Interestingly, TLR activation by viral or bacterial exposure in *early* pregnancy is more likely to result in complications than *late* pregnancy exposure (Allanson et al., 2010; Carpentier et al., 2011a, 2013; Hsiao and Patterson, 2011; Srinivas et al., 2006). Here, the acute effects of Poly(I:C) on immune-cell profiles differed between E12.5 and E14.5 (Figures 6 and S6).

Notably, changes in protein expression on mononuclear phagocytes and neutrophils were compartment and day dependent (Figure 6E). Extensive architectural changes between E12.5 and E14.5, with blood space expansion of the placenta being the most relevant (Coan et al., 2005; Coan et al., 2004), may offer some insight into the differential response to Poly(I:C) at early and late timepoints.

If the continual recruitment of immunomodulatory mononuclear phagocytes and neutrophils is required for normal placental homeostasis, we would anticipate that systemic immune perturbations would affect this recruitment and disrupt their enrichment and inferred functionality in placental tissue – a speculation that we hope to explore in future experiments. Additionally, to better understand how systemic activation can impact the placenta for the duration of gestation, future studies including longer intervals between activation and analysis are warranted.

Further investigation separating the maternal decidua from the fetal chorion is also warranted because our study preserved decidual attachment to the placenta and immune cell composition and phenotypic diversity is likely to differ in these two discrete microenvironments. An additional strategy to distinguish between maternal and fetal immune cells is needed to supplement the congenic mating strategy to confirm and fully characterize fetal immune contribution. Our analysis included a limited number of fetal cells and a small fraction could have been mischaracterized due to trogocytosis or non-specific antibody binding from the mechanical processing of tissue. Future studies could include allogeneic pregnancies to address maternal-fetal tolerance and to characterize any unique subpopulations that might only be present in an allogeneic context. Our characterization of myeloid subpopulations at the interface could be paired with single-cell transcriptional profiles to provide a richer overview of myeloid diversity. Future studies could also use our phenotypic profiles to detect and isolate mononuclear phagocyte and neutrophil subpopulations for further characterization of their development, identity, and function at the maternal-fetal interface.

In summary, our deep immune profiling of the tissue and endovascular compartments of the placenta and peripheral blood of 47 mice and ∼3 million cells across nine consecutive embryonic days enabled us to identify novel and noncanonical neutrophil and mononuclear phagocyte subpopulations linked to gestational immune remodeling. This resource provides a broad overview of compartmental immune dynamics in the placenta during pregnancy. These results suggest that innate immune cell diversity has been underappreciated, indicating the utility of high-dimensional approaches for tackling the complexities of the maternal-fetal interface.

Overall, the analytical framework, differentiation of the roles and composition of the placental tissue and endovascular compartments, and emphasis on the dynamics of (regulatory) innate immune populations provides a deeper foundation for pregnancy and its related pathologies.

## Acknowledgements

Thanks to Juliana Idoyaga, Virginia Winn, and Olivia Martinez for discussion and feedback; Michael Angelo and Joseph Campbell for critical review of the manuscript; Matthew Ball for computational support; Zinaida Good, David Glass, Erin McCaffrey, and Noah Greenwald for advice on data processing and visualization; Amy Moon, Jennifer Su, Aditya Asokan, Andrea Ocampo, Michelle Kielhold, Aditi Narayan, Anne Sommer, and Shelby Crants for experimental assistance; Irina Gurevich for technical assistance and resources; Michal C. Tal and Ying Y. Yiu for feedback and resources; and to the Stanford Immunology Graduate Program for training and support. A.R.M. was supported by the National Science Foundation Graduate Research Fellowship (DGE-1147470), the Stanford Diversifying Academia and Recruiting Excellence Doctoral Fellowship, and the Eunice Kennedy Shriver National Institute of Child Health and Human Development NIH-F31 Predoctoral Fellowship (5F31HD095569-03). N.V.G. was supported by the National Institute of Allergy and Infectious Diseases NIH-T32 Training Grant (5T32AI007290-35) and the Blavatnik Family Fellowship. T.D.P. was supported by NIH/NIMH 1R01MH108660-01, NIH/NIMH 1R01MH108659-01, Simons Foundation Research Grant 323220, NIH/NIMH R01MH09681501, and a SFARI Pilot Award. S.C.B. was supported by the DRCRF Fellowship (DRG-2017-09), the NIH 1DP2OD022550-01, 1R01AG056287-01, 1R01AG057915-01, R01AG068279, 1U24CA224309-01, 5U19AI116484-02, UH3 CA246633, U19 AG065156-01, and the Bill and Melinda Gates Foundation. Figures were created using BioRender (http://www.biorender.com) and Illustrator (Adobe).

## Author Contributions

A.R.M., N.V.G., S.C.B., and T.D.P. conceived and designed the study. A.R.M., N.V.G., K.A.P., O.R.M., and H.K. performed experiments. M.R. provided technical assistance with mouse experiments. B.C. performed statistical modeling. A.R.M. and N.V.G. performed data analysis and wrote the manuscript. S.C.B. and T.D.P. edited the manuscript. S.C.B. supervised the study and provided advice. S.C.B. and T.D.P. provided funding. All authors approved the final version of the manuscript.

## Code and data availability

All code and data will be made publicly available at the time of publication.

