## Supplementary material for "Gestationally-Dependent Immune Organization at the Maternal-Fetal Interface": SI and Methods

SUPPLEMENTAL TABLES and FIGURES

**Table S1. A summary of antibodies used for our mass cytometry analysis.** Related to Figure 1. The table shows the antibody target, antibody clone, the element and isotope the antibody was conjugated to, the final concentration of the metal-conjugated antibody used for protein detection, and the vendor from whom the antibodies were purchased.

| Antibody target | Clone | Element | Mass | Concentration (ug/ml) | Vendor |
|---|---|---|---|---|---|
| **Biotin** | 1D4-C5 | In | 113 | 2 | BioLegend |
| **CD45.2** | 104 | In | 115 | 4 | BioLegend |
| **Ter119** | TER-119 | La | 139 | 2 | BioLegend |
| **B220** | RA3-6B2 | Ce | 140 | 4 | BioLegend |
| **Ly-6G** | 1A8 | Pr | 141 | 2 | BioLegend |
| **CD11c** | N418 | Nd | 142 | 1 | BioLegend |
| **TCRβ** | H57-597 | Nd | 143 | 1 | BioLegend |
| **CD115 (CSF-1R)** | AFS98 | Nd | 144 | 2 | Fluidigm Sciences |
| **CD69** | H1.2F3 | Nd | 145 | 1 | BioLegend |
| **F4/80** | BM8 | Nd | 146 | 1 | BioLegend |
| **CD3** | 17A2 | Sm | 147 | 2 | BioLegend |
| **IgD** | 11-26c.2a | Nd | 148 | 2 | BioLegend |
| **CD19** | 6D5 | Sm | 149 | 1 | BioLegend |
| **CD25** | 3C7 | Nd | 150 | 4 | Fluidigm Sciences |
| **CD64** | X54-5/7.1 | Eu | 151 | 4 | BioLegend |
| **CD80** | 16-10A1 | Sm | 152 | 2 | BioLegend |
| **CD8** | 53-6.7 | Eu | 153 | 1 | BioLegend |
| **CD11b** | M1/70 | Sm | 154 | 1 | BioLegend |
| **CD40** | HM40-3 | Gd | 155 | 2 | BioLegend |
| **IgM** | RMM-1 | Gd | 156 | 1 | BioLegend |
| **CD117 (c-Kit)** | 2B8 | Gd | 157 | 0.5 | BioLegend |
| **TCRγδ** | GL3 | Tb | 159 | 2 | BioLegend |
| **CTLA-4 (CD152)** | UC10-4B9 | Dy | 161 | 2 | BioLegend |
| **Ly-6C** | HK1.4 | Dy | 162 | 2 | BioLegend |
| **CD194 (CCR4)** | 2G12 | Dy | 163 | 2 | BioLegend |
| **CD62L** | MEL-14 | Dy | 164 | 2 | BioLegend |
| **PD-L1 (CD274)** | 10F.9G2 | Ho | 165 | 2 | BioLegend |
| **FcεRI-α** | MAR-1 | Er | 166 | 0.5 | BioLegend |
| **CD335 (Nkp46)** | 29A1.4 | Er | 167 | 2 | BioLegend |
| **Siglec-F** | E50-2440 | Er | 168 | 1 | BDBiosciences |
| **CD49b** | DX5 | Er | 170 | 2 | BioLegend |
| **CD44** | IM7 | Yb | 171 | 1 | BioLegend |
| **CD4** | RM4-5 | Yb | 172 | 1 | BioLegend |
| **PD-1 (CD279)** | 29F.1A12 | Yb | 173 | 2 | BioLegend |
| **MHC-II** | M5/114.15.2 | Yb | 174 | 1 | BioLegend |
| **CD86** | GL-1 | Lu | 175 | 2 | BioLegend |
| **CD45.1** | A20 | Yb | 176 | 2 | BioLegend |
| **FoxP3** | FJK-16s | Gd | 158 | 4 | Fluidigm Sciences |
| **CD68** | FA-11 | Tm | 169 | 2 | BioLegend |
| **prpS6** | N7-548 | Bi | 209 | 2 | BD PhosphoFlow |

**Table S2. Microarray immune-associated genes.** Related to Figure 3B. The table shows the expression of immune-associated genes that are differentially expressed between E8.5 and 15.5 in the publicly available placental microarray data from Knox and Baker (2008).

**Table S3. Microarray vascular-associated genes.** Related to Figure 3B. The table shows the expression of vascular-associated genes that are differentially expressed between E8.5 and 15.5 in the publicly available placental microarray data from Knox and Baker (2008).

**Table S4. Generalized Estimating Equations coefficients and statistics for regressions of immune cell fractions over embryonic days.** Related to Figure 3C. Generalized Estimating Equations (GEE) was used to fit linear models and compare compartments within each cell type. PB was used as the reference.

| Coefficient | Estimate | Naive.S.E. | Naive.z | Robust.S.E. | Robust.z | P value | Cell type |
|---|---|---|---|---|---|---|---|
| **Intercept** | 2.66E-01 | 6.48E-02 | 4.11E+00 | 5.50E-02 | 4.84E+00 | 6.41E-07 | B cells |
| **Day** | 3.93E-03 | 4.38E-03 | 8.96E-01 | 3.59E-03 | 1.09E+00 | 1.37E-01 | B cells |
| **EV** | -2.14E-01 | 9.17E-02 | -2.33E+00 | 6.63E-02 | -3.22E+00 | 6.34E-04 | B cells |
| **TIS** | -2.26E-01 | 9.17E-02 | -2.46E+00 | 8.83E-02 | -2.56E+00 | 5.29E-03 | B cells |
| **Day:EV** | 3.81E-03 | 6.19E-03 | 6.15E-01 | 4.51E-03 | 8.44E-01 | 1.99E-01 | B cells |
| **Day:TIS** | -1.45E-03 | 6.19E-03 | -2.35E-01 | 6.21E-03 | -2.34E-01 | 4.07E-01 | B cells |
| **Intercept** | 2.55E-02 | 1.68E-02 | 1.52E+00 | 1.68E-02 | 1.52E+00 | 6.42E-02 | Basophils |
| **Day** | -7.89E-04 | 1.14E-03 | -6.91E-01 | 1.03E-03 | -7.63E-01 | 2.23E-01 | Basophils |
| **EV** | -9.31E-03 | 2.55E-02 | -3.65E-01 | 1.72E-02 | -5.42E-01 | 2.94E-01 | Basophils |
| **TIS** | 2.67E-03 | 2.37E-02 | 1.13E-01 | 2.46E-02 | 1.08E-01 | 4.57E-01 | Basophils |
| **Day:EV** | 4.37E-05 | 1.72E-03 | 2.55E-02 | 1.06E-03 | 4.13E-02 | 4.84E-01 | Basophils |
| **Day:TIS** | 1.49E-03 | 1.60E-03 | 9.28E-01 | 1.64E-03 | 9.07E-01 | 1.82E-01 | Basophils |
| **Intercept** | 8.86E-02 | 2.77E-02 | 3.20E+00 | 4.79E-02 | 1.85E+00 | 3.23E-02 | CD4+ T cells |
| **Day** | -9.08E-04 | 1.87E-03 | -4.86E-01 | 3.04E-03 | -2.99E-01 | 3.82E-01 | CD4+ T cells |
| **EV** | -1.20E-01 | 3.91E-02 | -3.06E+00 | 4.98E-02 | -2.40E+00 | 8.14E-03 | CD4+ T cells |
| **TIS** | -7.16E-02 | 3.91E-02 | -1.83E+00 | 5.24E-02 | -1.37E+00 | 8.58E-02 | CD4+ T cells |
| **Day:EV** | 5.30E-03 | 2.64E-03 | 2.01E+00 | 3.21E-03 | 1.65E+00 | 4.93E-02 | CD4+ T cells |
| **Day:TIS** | 1.48E-03 | 2.64E-03 | 5.61E-01 | 3.37E-03 | 4.39E-01 | 3.30E-01 | CD4+ T cells |
| **Intercept** | 8.33E-02 | 2.07E-02 | 4.03E+00 | 3.57E-02 | 2.33E+00 | 9.84E-03 | CD8+ T cells |
| **Day** | -1.14E-03 | 1.40E-03 | -8.16E-01 | 2.30E-03 | -4.96E-01 | 3.10E-01 | CD8+ T cells |
| **EV** | -1.09E-01 | 2.92E-02 | -3.72E+00 | 3.66E-02 | -2.97E+00 | 1.51E-03 | CD8+ T cells |
| **TIS** | -6.50E-02 | 2.92E-02 | -2.23E+00 | 4.04E-02 | -1.61E+00 | 5.37E-02 | CD8+ T cells |

| | | | | | | | |
|---|---|---|---|---|---|---|---|
| **Day:EV** | 4.48E-03 | 1.97E-03 | 2.27E+00 | 2.36E-03 | 1.90E+00 | 2.85E-02 | CD8+ T cells |
| **Day:TIS** | 1.18E-03 | 1.97E-03 | 5.99E-01 | 2.67E-03 | 4.42E-01 | 3.29E-01 | CD8+ T cells |
| **Intercept** | 3.17E-02 | 1.36E-02 | 2.33E+00 | 1.95E-02 | 1.62E+00 | 5.24E-02 | Eosinophils |
| **Day** | -1.60E-03 | 9.22E-04 | -1.74E+00 | 1.17E-03 | -1.36E+00 | 8.64E-02 | Eosinophils |
| **EV** | -4.06E-03 | 1.91E-02 | -2.12E-01 | 2.26E-02 | -1.80E-01 | 4.29E-01 | Eosinophils |
| **TIS** | 8.14E-03 | 1.91E-02 | 4.25E-01 | 2.31E-02 | 3.53E-01 | 3.62E-01 | Eosinophils |
| **Day:EV** | 5.33E-04 | 1.30E-03 | 4.11E-01 | 1.36E-03 | 3.93E-01 | 3.47E-01 | Eosinophils |
| **Day:TIS** | 5.59E-04 | 1.30E-03 | 4.31E-01 | 1.46E-03 | 3.84E-01 | 3.51E-01 | Eosinophils |
| **Intercept** | 1.80E-01 | 7.10E-02 | 2.53E+00 | 4.68E-02 | 3.85E+00 | 6.00E-05 | Monocytes |
| **Day** | -4.48E-03 | 4.80E-03 | -9.34E-01 | 3.04E-03 | -1.47E+00 | 7.06E-02 | Monocytes |
| **EV** | 9.79E-01 | 1.00E-01 | 9.75E+00 | 6.72E-02 | 1.46E+01 | 2.02E-48 | Monocytes |
| **TIS** | 3.72E-01 | 1.00E-01 | 3.70E+00 | 8.18E-02 | 4.54E+00 | 2.80E-06 | Monocytes |
| **Day:EV** | -3.91E-02 | 6.78E-03 | -5.77E+00 | 4.63E-03 | -8.44E+00 | 1.55E-17 | Monocytes |
| **Day:TIS** | -6.67E-03 | 6.78E-03 | -9.83E-01 | 5.77E-03 | -1.15E+00 | 1.24E-01 | Monocytes |
| **Intercept** | 2.46E-01 | 5.42E-02 | 4.54E+00 | 7.94E-02 | 3.10E+00 | 9.77E-04 | NK cells |
| **Day** | 3.90E-04 | 3.66E-03 | 1.06E-01 | 5.30E-03 | 7.36E-02 | 4.71E-01 | NK cells |
| **EV** | -2.46E-01 | 7.67E-02 | -3.21E+00 | 8.06E-02 | -3.05E+00 | 1.14E-03 | NK cells |
| **TIS** | -8.02E-02 | 7.67E-02 | -1.05E+00 | 8.63E-02 | -9.29E-01 | 1.76E-01 | NK cells |
| **Day:EV** | 2.63E-03 | 5.18E-03 | 5.08E-01 | 5.38E-03 | 4.90E-01 | 3.12E-01 | NK cells |
| **Day:TIS** | -5.82E-03 | 5.18E-03 | -1.12E+00 | 5.77E-03 | -1.01E+00 | 1.56E-01 | NK cells |
| **Intercept** | 7.59E-02 | 8.43E-02 | 9.01E-01 | 3.75E-02 | 2.02E+00 | 2.15E-02 | Neutrophils |
| **Day** | 4.86E-03 | 5.69E-03 | 8.54E-01 | 2.58E-03 | 1.88E+00 | 2.98E-02 | Neutrophils |
| **EV** | -2.70E-01 | 1.19E-01 | -2.26E+00 | 6.67E-02 | -4.04E+00 | 2.63E-05 | Neutrophils |
| **TIS** | 6.28E-02 | 1.19E-01 | 5.26E-01 | 9.32E-02 | 6.73E-01 | 2.50E-01 | Neutrophils |
| **Day:EV** | 2.18E-02 | 8.05E-03 | 2.71E+00 | 4.67E-03 | 4.67E+00 | 1.53E-06 | Neutrophils |
| **Day:TIS** | 8.97E-03 | 8.05E-03 | 1.11E+00 | 6.77E-03 | 1.32E+00 | 9.26E-02 | Neutrophils |

**Table S5. Generalized Estimating Equations coefficients and statistics for regressions of protein medians in MP and neutrophils over embryonic days.** Related to Figure 3G. GEE was used to fit linear models per protein and cell type to compare compartments. PB was used as the reference.

| Coefficient | Estimate | Naive.S.E. | Naive.z | Robust.S.E. | Robust.z | P value | Cell/protein |
|---|---|---|---|---|---|---|---|
| **Intercept** | 7.92E-01 | 5.71E-01 | 1.39E+00 | 3.58E-01 | 2.21E+00 | 1.34E-02 | MP/PD-L1 |
| **Day** | -3.01E-02 | 3.85E-02 | -7.80E-01 | 2.28E-02 | -1.32E+00 | 9.32E-02 | MP/PD-L1 |
| **EV** | 1.84E+00 | 8.07E-01 | 2.28E+00 | 9.41E-01 | 1.95E+00 | 2.54E-02 | MP/PD-L1 |
| **TIS** | 3.65E+00 | 8.07E-01 | 4.52E+00 | 6.28E-01 | 5.81E+00 | 3.20E-09 | MP/PD-L1 |
| **Day:EV** | -1.17E-01 | 5.45E-02 | -2.15E+00 | 5.85E-02 | -2.01E+00 | 2.25E-02 | MP/PD-L1 |
| **Day:TIS** | -1.76E-01 | 5.45E-02 | -3.22E+00 | 4.13E-02 | -4.26E+00 | 1.03E-05 | MP/PD-L1 |
| **Intercept** | 3.49E-01 | 1.45E-01 | 2.41E+00 | 1.19E-01 | 2.94E+00 | 1.63E-03 | Neutrophils/PD-L1 |
| **Day** | -1.28E-02 | 9.78E-03 | -1.30E+00 | 7.75E-03 | -1.65E+00 | 5.00E-02 | Neutrophils/PD-L1 |
| **EV** | -7.80E-02 | 2.05E-01 | -3.81E-01 | 1.77E-01 | -4.40E-01 | 3.30E-01 | Neutrophils/PD-L1 |
| **TIS** | 3.47E-01 | 2.05E-01 | 1.69E+00 | 1.93E-01 | 1.80E+00 | 3.59E-02 | Neutrophils/PD-L1 |
| **Day:EV** | 3.70E-03 | 1.38E-02 | 2.68E-01 | 1.08E-02 | 3.42E-01 | 3.66E-01 | Neutrophils/PD-L1 |
| **Day:TIS** | -1.33E-02 | 1.38E-02 | -9.63E-01 | 1.27E-02 | -1.05E+00 | 1.47E-01 | Neutrophils/PD-L1 |
| **Intercept** | 4.55E+00 | 6.85E-01 | 6.64E+00 | 9.90E-01 | 4.59E+00 | 2.20E-06 | MP/Ly-6C |
| **Day** | 8.76E-03 | 4.63E-02 | 1.89E-01 | 6.59E-02 | 1.33E-01 | 4.47E-01 | MP/Ly-6C |
| **EV** | -9.56E-01 | 9.69E-01 | -9.87E-01 | 9.73E-01 | -9.83E-01 | 1.63E-01 | MP/Ly-6C |
| **TIS** | 1.12E+00 | 9.69E-01 | 1.16E+00 | 9.55E-01 | 1.18E+00 | 1.20E-01 | MP/Ly-6C |
| **Day:EV** | 7.25E-02 | 6.54E-02 | 1.11E+00 | 6.62E-02 | 1.10E+00 | 1.36E-01 | MP/Ly-6C |
| **Day:TIS** | -1.10E-01 | 6.54E-02 | -1.69E+00 | 6.93E-02 | -1.59E+00 | 5.55E-02 | MP/Ly-6C |
| **Intercept** | 2.97E+00 | 2.67E-01 | 1.11E+01 | 3.89E-01 | 7.64E+00 | 1.12E-14 | Neutrophils/Ly-6G |
| **Day** | -4.26E-03 | 1.80E-02 | -2.36E-01 | 2.50E-02 | -1.70E-01 | 4.32E-01 | Neutrophils/Ly-6G |
| **EV** | -1.17E+00 | 3.77E-01 | -3.09E+00 | 3.35E-01 | -3.48E+00 | 2.54E-04 | Neutrophils/Ly-6G |
| **TIS** | -1.31E+00 | 3.77E-01 | -3.47E+00 | 3.42E-01 | -3.83E+00 | 6.36E-05 | Neutrophils/Ly-6G |
| **Day:EV** | 6.39E-02 | 2.55E-02 | 2.51E+00 | 2.17E-02 | 2.95E+00 | 1.58E-03 | Neutrophils/Ly-6G |
| **Day:TIS** | 6.30E-02 | 2.55E-02 | 2.47E+00 | 2.31E-02 | 2.72E+00 | 3.22E-03 | Neutrophils/Ly-6G |

**Table S6. Generalized Estimating Equations coefficients and statistics for regressions of immune fractions in MP subsets over embryonic days.** Related to Figure 4E and S4G. GEE was used to fit linear models and compare fraction of immune cells in compartments within each MP subset. PB was used as the reference.

| Coefficient | Estimate | Naive.S.E. | Naive.z | Robust.S.E. | Robust.z | P value | Cell type |
|---|---|---|---|---|---|---|---|
| **Intercept** | 6.69E-02 | 3.97E-02 | 1.68E+00 | 3.28E-02 | 2.04E+00 | 2.06E-02 | Ly-6Chi |
| **Day** | -4.63E-04 | 2.68E-03 | -1.73E-01 | 2.04E-03 | -2.27E-01 | 4.10E-01 | Ly-6Chi |
| **EV** | 1.02E-01 | 5.61E-02 | 1.82E+00 | 5.88E-02 | 1.74E+00 | 4.11E-02 | Ly-6Chi |
| **TIS** | 3.41E-02 | 5.61E-02 | 6.08E-01 | 5.01E-02 | 6.81E-01 | 2.48E-01 | Ly-6Chi |
| **Day:EV** | 3.34E-03 | 3.79E-03 | 8.82E-01 | 3.86E-03 | 8.66E-01 | 1.93E-01 | Ly-6Chi |
| **Day:TIS** | 4.79E-04 | 3.79E-03 | 1.26E-01 | 3.30E-03 | 1.45E-01 | 4.42E-01 | Ly-6Chi |
| **Intercept** | 3.98E-02 | 1.84E-02 | 2.17E+00 | 1.09E-02 | 3.64E+00 | 1.34E-04 | Patrolling |
| **Day** | -1.29E-03 | 1.24E-03 | -1.04E+00 | 7.14E-04 | -1.81E+00 | 3.55E-02 | Patrolling |
| **EV** | 1.73E-01 | 2.60E-02 | 6.65E+00 | 2.96E-02 | 5.85E+00 | 2.50E-09 | Patrolling |
| **TIS** | 9.81E-03 | 2.60E-02 | 3.77E-01 | 1.60E-02 | 6.13E-01 | 2.70E-01 | Patrolling |
| **Day:EV** | -9.79E-03 | 1.76E-03 | -5.58E+00 | 1.91E-03 | -5.11E+00 | 1.57E-07 | Patrolling |
| **Day:TIS** | -5.33E-04 | 1.76E-03 | -3.03E-01 | 1.05E-03 | -5.05E-01 | 3.07E-01 | Patrolling |
| **Intercept** | 1.05E-02 | 3.19E-02 | 3.30E-01 | 1.43E-03 | 7.37E+00 | 8.71E-14 | Phagocytic |
| **Day** | -4.30E-04 | 2.11E-03 | -2.04E-01 | 9.19E-05 | -4.68E+00 | 1.42E-06 | Phagocytic |
| **EV** | 5.79E-02 | 3.95E-02 | 1.47E+00 | 2.16E-02 | 2.68E+00 | 3.63E-03 | Phagocytic |
| **TIS** | 1.55E-02 | 3.95E-02 | 3.92E-01 | 2.86E-02 | 5.41E-01 | 2.94E-01 | Phagocytic |
| **Day:EV** | -2.57E-03 | 2.63E-03 | -9.76E-01 | 1.35E-03 | -1.91E+00 | 2.84E-02 | Phagocytic |
| **Day:TIS** | 3.21E-03 | 2.63E-03 | 1.22E+00 | 2.02E-03 | 1.59E+00 | 5.62E-02 | Phagocytic |
| **Intercept** | -2.30E-04 | 7.59E-02 | -3.03E-03 | 1.51E-03 | -1.53E-01 | 4.39E-01 | MoDC |
| **Day** | 2.13E-04 | 5.73E-03 | 3.72E-02 | 1.31E-04 | 1.63E+00 | 5.20E-02 | MoDC |

| | | | | | | | |
|---|---|---|---|---|---|---|---|
| **EV** | 2.23E-01 | 8.31E-02 | 2.68E+00 | 5.44E-02 | 4.10E+00 | 2.06E-05 | MoDC |
| **TIS** | 1.83E-01 | 8.31E-02 | 2.20E+00 | 2.18E-02 | 8.36E+00 | 3.01E-17 | MoDC |
| **Day:EV** | -1.24E-02 | 6.17E-03 | -2.01E+00 | 3.39E-03 | -3.66E+00 | 1.29E-04 | MoDC |
| **Day:TIS** | -8.41E-03 | 6.17E-03 | -1.36E+00 | 1.49E-03 | -5.63E+00 | 8.93E-09 | MoDC |
| **Intercept** | 7.84E-03 | 1.12E-02 | 7.01E-01 | 4.87E-03 | 1.61E+00 | 5.36E-02 | DC |
| **Day** | -1.98E-04 | 7.80E-04 | -2.54E-01 | 3.24E-04 | -6.11E-01 | 2.71E-01 | DC |
| **EV** | 2.42E-02 | 1.53E-02 | 1.58E+00 | 1.32E-02 | 1.84E+00 | 3.32E-02 | DC |
| **TIS** | 8.80E-03 | 1.53E-02 | 5.76E-01 | 9.62E-03 | 9.15E-01 | 1.80E-01 | DC |
| **Day:EV** | 2.77E-05 | 1.05E-03 | 2.64E-02 | 8.49E-04 | 3.26E-02 | 4.87E-01 | DC |
| **Day:TIS** | 4.00E-04 | 1.05E-03 | 3.81E-01 | 6.84E-04 | 5.84E-01 | 2.80E-01 | DC |
| **Intercept** | 6.01E-02 | 2.68E-02 | 2.25E+00 | 1.00E-02 | 5.99E+00 | 1.07E-09 | Presenting |
| **Day** | -2.78E-03 | 1.81E-03 | -1.54E+00 | 6.34E-04 | -4.39E+00 | 5.73E-06 | Presenting |
| **EV** | 2.60E-01 | 3.78E-02 | 6.87E+00 | 3.30E-02 | 7.86E+00 | 1.86E-15 | Presenting |
| **TIS** | 1.71E-02 | 3.78E-02 | 4.52E-01 | 2.54E-02 | 6.73E-01 | 2.50E-01 | Presenting |
| **Day:EV** | -9.92E-03 | 2.56E-03 | -3.88E+00 | 2.26E-03 | -4.39E+00 | 5.54E-06 | Presenting |
| **Day:TIS** | 1.48E-03 | 2.56E-03 | 5.81E-01 | 1.81E-03 | 8.19E-01 | 2.06E-01 | Presenting |
| **Intercept** | 4.58E-03 | 1.84E-02 | 2.49E-01 | 1.94E-03 | 2.36E+00 | 9.05E-03 | Proliferating |
| **Day** | -1.58E-04 | 1.29E-03 | -1.23E-01 | 1.25E-04 | -1.27E+00 | 1.02E-01 | Proliferating |
| **EV** | 4.80E-02 | 2.04E-02 | 2.35E+00 | 7.05E-03 | 6.81E+00 | 4.75E-12 | Proliferating |
| **TIS** | 5.72E-02 | 2.03E-02 | 2.82E+00 | 1.08E-02 | 5.29E+00 | 6.20E-08 | Proliferating |
| **Day:EV** | -2.86E-03 | 1.43E-03 | -2.00E+00 | 4.55E-04 | -6.28E+00 | 1.71E-10 | Proliferating |
| **Day:TIS** | -3.11E-03 | 1.42E-03 | -2.19E+00 | 6.85E-04 | -4.54E+00 | 2.77E-06 | Proliferating |
| **Intercept** | 5.55E-03 | 1.11E-01 | 4.99E-02 | 9.78E-02 | 5.68E-02 | 4.77E-01 | F4/80hi |
| **Day** | 5.95E-03 | 7.44E-03 | 8.00E-01 | 7.33E-03 | 8.12E-01 | 2.08E-01 | F4/80hi |

.

**Table S7. Generalized Estimating Equations coefficients and statistics for regressions of PD-L1+ MP fractions in PD-L1+ MP subsets over embryonic days.** Related to Figure 4F. GEE was used to fit linear models and compare fraction of PD-L1+ MPs in compartments within each PD-L1+ MP subset. PB was used as the reference.

| Coefficient | Estimate | Naive.S.E. | Naive.z | Robust.S.E. | Robust.z | P value | Cell type |
|---|---|---|---|---|---|---|---|
| **Intercept** | 8.05E-01 | 1.41E-01 | 5.72E+00 | 1.30E-01 | 6.19E+00 | 3.06E-10 | Patrolling |
| **Day** | 5.23E-03 | 9.50E-03 | 5.51E-01 | 8.42E-03 | 6.21E-01 | 2.67E-01 | Patrolling |
| **EV** | -4.08E-01 | 1.99E-01 | -2.05E+00 | 2.13E-01 | -1.92E+00 | 2.75E-02 | Patrolling |
| **TIS** | -6.24E-01 | 1.99E-01 | -3.14E+00 | 1.60E-01 | -3.91E+00 | 4.64E-05 | Patrolling |
| **Day:EV** | -2.53E-03 | 1.34E-02 | -1.89E-01 | 1.38E-02 | -1.84E-01 | 4.27E-01 | Patrolling |
| **Day:TIS** | -6.35E-03 | 1.34E-02 | -4.73E-01 | 1.07E-02 | -5.93E-01 | 2.77E-01 | Patrolling |
| **Intercept** | 3.28E-01 | 1.33E-01 | 2.46E+00 | 6.66E-02 | 4.92E+00 | 4.32E-07 | Phagocytic |
| **Day** | -1.14E-02 | 8.81E-03 | -1.30E+00 | 4.31E-03 | -2.65E+00 | 4.00E-03 | Phagocytic |
| **EV** | -3.40E-01 | 1.65E-01 | -2.06E+00 | 9.87E-02 | -3.45E+00 | 2.82E-04 | Phagocytic |
| **TIS** | -4.65E-01 | 1.65E-01 | -2.82E+00 | 1.24E-01 | -3.75E+00 | 8.97E-05 | Phagocytic |
| **Day:EV** | 2.80E-02 | 1.10E-02 | 2.55E+00 | 6.41E-03 | 4.37E+00 | 6.26E-06 | Phagocytic |
| **Day:TIS** | 5.12E-02 | 1.10E-02 | 4.66E+00 | 8.86E-03 | 5.78E+00 | 3.69E-09 | Phagocytic |
| **Intercept** | -1.44E-01 | 2.26E-01 | -6.35E-01 | 9.58E-02 | -1.50E+00 | 6.67E-02 | MoDC |
| **Day** | 1.80E-02 | 1.71E-02 | 1.05E+00 | 7.76E-03 | 2.31E+00 | 1.03E-02 | MoDC |
| **EV** | 7.60E-01 | 2.48E-01 | 3.07E+00 | 1.62E-01 | 4.68E+00 | 1.44E-06 | MoDC |
| **TIS** | 1.10E+00 | 2.48E-01 | 4.44E+00 | 1.20E-01 | 9.19E+00 | 2.02E-20 | MoDC |
| **Day:EV** | -3.73E-02 | 1.84E-02 | -2.03E+00 | 1.18E-02 | -3.17E+00 | 7.63E-04 | MoDC |
| **Day:TIS** | -5.66E-02 | 1.84E-02 | -3.08E+00 | 9.27E-03 | -6.11E+00 | 5.04E-10 | MoDC |

**Table S8. Generalized Estimating Equations coefficients and statistics for regressions of fractions out of MPs in MP subsets over embryonic days.** Related to Figure 4 and S4H. GEE was used to fit linear models and compare fraction out of MPs in compartments within each MP subset. PB was used as the reference.

| Coefficient | Estimate | Naive.S.E. | Naive.z | Robust.S.E. | Robust.z | P value | Cell type |
|---|---|---|---|---|---|---|---|
| **Intercept** | 6.69E-02 | 3.97E-02 | 1.68E+00 | 3.28E-02 | 2.04E+00 | 2.06E-02 | Ly-6Chi |
| **Day** | -4.63E-04 | 2.68E-03 | -1.73E-01 | 2.04E-03 | -2.27E-01 | 4.10E-01 | Ly-6Chi |
| **EV** | 1.02E-01 | 5.61E-02 | 1.82E+00 | 5.88E-02 | 1.74E+00 | 4.11E-02 | Ly-6Chi |
| **TIS** | 3.41E-02 | 5.61E-02 | 6.08E-01 | 5.01E-02 | 6.81E-01 | 2.48E-01 | Ly-6Chi |
| **Day:EV** | 3.34E-03 | 3.79E-03 | 8.82E-01 | 3.86E-03 | 8.66E-01 | 1.93E-01 | Ly-6Chi |
| **Day:TIS** | 4.79E-04 | 3.79E-03 | 1.26E-01 | 3.30E-03 | 1.45E-01 | 4.42E-01 | Ly-6Chi |
| **Intercept** | 3.98E-02 | 1.84E-02 | 2.17E+00 | 1.09E-02 | 3.64E+00 | 1.34E-04 | Patrolling |
| **Day** | -1.29E-03 | 1.24E-03 | -1.04E+00 | 7.14E-04 | -1.81E+00 | 3.55E-02 | Patrolling |
| **EV** | 1.73E-01 | 2.60E-02 | 6.65E+00 | 2.96E-02 | 5.85E+00 | 2.50E-09 | Patrolling |
| **TIS** | 9.81E-03 | 2.60E-02 | 3.77E-01 | 1.60E-02 | 6.13E-01 | 2.70E-01 | Patrolling |
| **Day:EV** | -9.79E-03 | 1.76E-03 | -5.58E+00 | 1.91E-03 | -5.11E+00 | 1.57E-07 | Patrolling |
| **Day:TIS** | -5.33E-04 | 1.76E-03 | -3.03E-01 | 1.05E-03 | -5.05E-01 | 3.07E-01 | Patrolling |
| **Intercept** | 1.05E-02 | 3.19E-02 | 3.30E-01 | 1.43E-03 | 7.37E+00 | 8.71E-14 | Phagocytic |
| **Day** | -4.30E-04 | 2.11E-03 | -2.04E-01 | 9.19E-05 | -4.68E+00 | 1.42E-06 | Phagocytic |
| **EV** | 5.79E-02 | 3.95E-02 | 1.47E+00 | 2.16E-02 | 2.68E+00 | 3.63E-03 | Phagocytic |
| **TIS** | 1.55E-02 | 3.95E-02 | 3.92E-01 | 2.86E-02 | 5.41E-01 | 2.94E-01 | Phagocytic |
| **Day:EV** | -2.57E-03 | 2.63E-03 | -9.76E-01 | 1.35E-03 | -1.91E+00 | 2.84E-02 | Phagocytic |
| **Day:TIS** | 3.21E-03 | 2.63E-03 | 1.22E+00 | 2.02E-03 | 1.59E+00 | 5.62E-02 | Phagocytic |
| **Intercept** | -2.30E-04 | 7.59E-02 | -3.03E-03 | 1.51E-03 | -1.53E-01 | 4.39E-01 | MoDC |
| **Day** | 2.13E-04 | 5.73E-03 | 3.72E-02 | 1.31E-04 | 1.63E+00 | 5.20E-02 | MoDC |
| **EV** | 2.23E-01 | 8.31E-02 | 2.68E+00 | 5.44E-02 | 4.10E+00 | 2.06E-05 | MoDC |

| | | | | | | | |
|---|---|---|---|---|---|---|---|
| **TIS** | 1.83E-01 | 8.31E-02 | 2.20E+00 | 2.18E-02 | 8.36E+00 | 3.01E-17 | MoDC |
| **Day:EV** | -1.24E-02 | 6.17E-03 | -2.01E+00 | 3.39E-03 | -3.66E+00 | 1.29E-04 | MoDC |
| **Day:TIS** | -8.41E-03 | 6.17E-03 | -1.36E+00 | 1.49E-03 | -5.63E+00 | 8.93E-09 | MoDC |
| **Intercept** | 7.84E-03 | 1.12E-02 | 7.01E-01 | 4.87E-03 | 1.61E+00 | 5.36E-02 | DC |
| **Day** | -1.98E-04 | 7.80E-04 | -2.54E-01 | 3.24E-04 | -6.11E-01 | 2.71E-01 | DC |
| **EV** | 2.42E-02 | 1.53E-02 | 1.58E+00 | 1.32E-02 | 1.84E+00 | 3.32E-02 | DC |
| **TIS** | 8.80E-03 | 1.53E-02 | 5.76E-01 | 9.62E-03 | 9.15E-01 | 1.80E-01 | DC |
| **Day:EV** | 2.77E-05 | 1.05E-03 | 2.64E-02 | 8.49E-04 | 3.26E-02 | 4.87E-01 | DC |
| **Day:TIS** | 4.00E-04 | 1.05E-03 | 3.81E-01 | 6.84E-04 | 5.84E-01 | 2.80E-01 | DC |
| **Intercept** | 6.01E-02 | 2.68E-02 | 2.25E+00 | 1.00E-02 | 5.99E+00 | 1.07E-09 | Presenting |
| **Day** | -2.78E-03 | 1.81E-03 | -1.54E+00 | 6.34E-04 | -4.39E+00 | 5.73E-06 | Presenting |
| **EV** | 2.60E-01 | 3.78E-02 | 6.87E+00 | 3.30E-02 | 7.86E+00 | 1.86E-15 | Presenting |
| **TIS** | 1.71E-02 | 3.78E-02 | 4.52E-01 | 2.54E-02 | 6.73E-01 | 2.50E-01 | Presenting |
| **Day:EV** | -9.92E-03 | 2.56E-03 | -3.88E+00 | 2.26E-03 | -4.39E+00 | 5.54E-06 | Presenting |
| **Day:TIS** | 1.48E-03 | 2.56E-03 | 5.81E-01 | 1.81E-03 | 8.19E-01 | 2.06E-01 | Presenting |
| **Intercept** | 4.58E-03 | 1.84E-02 | 2.49E-01 | 1.94E-03 | 2.36E+00 | 9.05E-03 | Proliferating |
| **Day** | -1.58E-04 | 1.29E-03 | -1.23E-01 | 1.25E-04 | -1.27E+00 | 1.02E-01 | Proliferating |
| **EV** | 4.80E-02 | 2.04E-02 | 2.35E+00 | 7.05E-03 | 6.81E+00 | 4.75E-12 | Proliferating |
| **TIS** | 5.72E-02 | 2.03E-02 | 2.82E+00 | 1.08E-02 | 5.29E+00 | 6.20E-08 | Proliferating |
| **Day:EV** | -2.86E-03 | 1.43E-03 | -2.00E+00 | 4.55E-04 | -6.28E+00 | 1.71E-10 | Proliferating |
| **Day:TIS** | -3.11E-03 | 1.42E-03 | -2.19E+00 | 6.85E-04 | -4.54E+00 | 2.77E-06 | Proliferating |
| **Intercept** | 5.55E-03 | 1.11E-01 | 4.99E-02 | 9.78E-02 | 5.68E-02 | 4.77E-01 | F4/80hi |
| **Day** | 5.95E-03 | 7.44E-03 | 8.00E-01 | 7.33E-03 | 8.12E-01 | 2.08E-01 | F4/80hi |

**Table S9. Generalized Estimating Equations coefficients and statistics for regressions of immune fractions in neutrophil subsets over embryonic days.** Related to Figure 5F. GEE was used to fit linear models to all but immunosuppressive neutrophil subset to compare fraction of immune cells in compartments within each neutrophil subset. Immunosuppressive subset was fitted with a quadratic model. PB was used as the reference.

| Coefficient | Estimate | Naive.S.E. | Naive.z | Robust.S.E. | Robust.z | P value | Cell type |
|---|---|---|---|---|---|---|---|
| **Intercept** | 2.83E-03 | 8.35E-03 | 3.38E-01 | 4.80E-03 | 5.89E-01 | 2.78E-01 | CD80 |
| **Day** | 6.48E-04 | 5.68E-04 | 1.14E+00 | 3.00E-04 | 2.16E+00 | 1.55E-02 | CD80 |
| **EV** | -1.93E-02 | 1.21E-02 | -1.60E+00 | 6.56E-03 | -2.94E+00 | 1.62E-03 | CD80 |
| **TIS** | -2.39E-02 | 1.17E-02 | -2.04E+00 | 1.03E-02 | -2.31E+00 | 1.05E-02 | CD80 |
| **Day:EV** | 1.34E-03 | 8.14E-04 | 1.65E+00 | 4.27E-04 | 3.14E+00 | 8.31E-04 | CD80 |
| **Day:TIS** | 1.89E-03 | 7.94E-04 | 2.39E+00 | 7.54E-04 | 2.51E+00 | 6.04E-03 | CD80 |
| **Intercept** | 1.34E-02 | 3.93E-02 | 3.40E-01 | 2.19E-02 | 6.11E-01 | 2.71E-01 | Conventional |
| **Day** | 2.81E-03 | 2.66E-03 | 1.06E+00 | 1.52E-03 | 1.84E+00 | 3.26E-02 | Conventional |
| **EV** | -7.82E-02 | 5.56E-02 | -1.41E+00 | 3.92E-02 | -2.00E+00 | 2.29E-02 | Conventional |
| **TIS** | 2.87E-02 | 5.56E-02 | 5.16E-01 | 3.16E-02 | 9.08E-01 | 1.82E-01 | Conventional |
| **Day:EV** | 7.16E-03 | 3.76E-03 | 1.91E+00 | 2.75E-03 | 2.60E+00 | 4.60E-03 | Conventional |
| **Day:TIS** | 7.74E-04 | 3.76E-03 | 2.06E-01 | 2.20E-03 | 3.51E-01 | 3.63E-01 | Conventional |
| **Intercept** | 2.77E-02 | 3.04E-02 | 9.10E-01 | 2.04E-02 | 1.36E+00 | 8.74E-02 | Presenting |
| **Day** | 2.09E-03 | 2.05E-03 | 1.02E+00 | 1.36E-03 | 1.54E+00 | 6.16E-02 | Presenting |
| **EV** | -1.56E-01 | 4.30E-02 | -3.62E+00 | 2.68E-02 | -5.81E+00 | 3.17E-09 | Presenting |
| **TIS** | -1.04E-01 | 4.30E-02 | -2.42E+00 | 3.94E-02 | -2.65E+00 | 4.08E-03 | Presenting |
| **Day:EV** | 1.18E-02 | 2.90E-03 | 4.07E+00 | 1.85E-03 | 6.39E+00 | 8.57E-11 | Presenting |
| **Day:TIS** | 9.11E-03 | 2.90E-03 | 3.14E+00 | 2.92E-03 | 3.12E+00 | 8.99E-04 | Presenting |
| **Intercept** | 1.00E-02 | 1.05E-02 | 9.56E-01 | 6.92E-03 | 1.45E+00 | 7.38E-02 | Proliferating |
| **Day** | -1.16E-04 | 7.01E-04 | -1.66E-01 | 4.39E-04 | -2.65E-01 | 3.95E-01 | Proliferating |
| **EV** | 1.17E-02 | 1.49E-02 | 7.84E-01 | 8.93E-03 | 1.31E+00 | 9.56E-02 | Proliferating |

| | | | | | | | |
|---|---|---|---|---|---|---|---|
| **TIS** | 9.64E-02 | 1.59E-02 | 6.07E+00 | 1.53E-02 | 6.29E+00 | 1.59E-10 | Proliferating |
| **Day:EV** | -8.77E-04 | 1.02E-03 | -8.62E-01 | 5.77E-04 | -1.52E+00 | 6.43E-02 | Proliferating |
| **Day:TIS** | -5.30E-03 | 1.11E-03 | -4.78E+00 | 1.01E-03 | -5.27E+00 | 6.68E-08 | Proliferating |
| **Intercept** | 1.33E-02 | 1.25E-02 | 1.07E+00 | 7.80E-03 | 1.71E+00 | 4.37E-02 | Immunosuppressive |
| **Day** | -4.71E-04 | 8.56E-04 | -5.50E-01 | 4.97E-04 | -9.47E-01 | 1.72E-01 | Immunosuppressive |
| **EV** | -3.19E-03 | 1.86E-02 | -1.72E-01 | 9.10E-03 | -3.51E-01 | 3.63E-01 | Immunosuppressive |
| **TIS** | -4.92E-01 | 7.36E-02 | -6.69E+00 | 1.03E-01 | -4.79E+00 | 8.49E-07 | Immunosuppressive |
| **Day:EV** | 3.92E-04 | 1.26E-03 | 3.11E-01 | 5.76E-04 | 6.80E-01 | 2.48E-01 | Immunosuppressive |
| **Day:TIS** | 8.02E-02 | 1.03E-02 | 7.80E+00 | 1.40E-02 | 5.75E+00 | 4.45E-09 | Immunosuppressive |
| **Day-sq:TIS** | -2.84E-03 | 3.53E-04 | -8.07E+00 | 4.59E-04 | -6.20E+00 | 2.83E-10 | Immunosuppressive |

**Table S10. Generalized Estimating Equations coefficients and statistics for regressions of fractions out of neutrophils in neutrophil subsets over embryonic days.** Related to Figure 5G. GEE was used to fit linear models to all but immunosuppressive neutrophil subset to compare fraction out of neutrophils in compartments within each neutrophil subset. Immunosuppressive subset was fitted with a quadratic model. PB was used as the reference.

| Coefficient | Estimate | Naive.S.E. | Naive.z | Robust.S.E. | Robust.z | P value | Cell type |
|---|---|---|---|---|---|---|---|
| **Intercept** | 4.88E-02 | 3.22E-02 | 1.52E+00 | 3.50E-02 | 1.39E+00 | 8.18E-02 | CD80 Neu |
| **Day** | 2.35E-03 | 2.19E-03 | 1.07E+00 | 2.24E-03 | 1.05E+00 | 1.47E-01 | CD80 Neu |
| **EV** | 5.05E-04 | 4.65E-02 | 1.08E-02 | 4.67E-02 | 1.08E-02 | 4.96E-01 | CD80 Neu |
| **TIS** | -9.49E-02 | 4.51E-02 | -2.11E+00 | 4.08E-02 | -2.33E+00 | 9.93E-03 | CD80 Neu |
| **Day:EV** | -1.16E-03 | 3.14E-03 | -3.69E-01 | 3.01E-03 | -3.85E-01 | 3.50E-01 | CD80 Neu |
| **Day:TIS** | 4.34E-03 | 3.06E-03 | 1.42E+00 | 2.70E-03 | 1.61E+00 | 5.39E-02 | CD80 Neu |
| **Intercept** | 2.54E-01 | 8.77E-02 | 2.89E+00 | 9.10E-02 | 2.79E+00 | 2.64E-03 | Conventional |
| **Day** | 7.11E-03 | 5.92E-03 | 1.20E+00 | 5.78E-03 | 1.23E+00 | 1.10E-01 | Conventional |
| **EV** | 1.07E-01 | 1.24E-01 | 8.64E-01 | 1.28E-01 | 8.36E-01 | 2.02E-01 | Conventional |
| **TIS** | 3.76E-02 | 1.24E-01 | 3.03E-01 | 1.04E-01 | 3.60E-01 | 3.59E-01 | Conventional |
| **Day:EV** | -4.32E-03 | 8.38E-03 | -5.16E-01 | 8.25E-03 | -5.24E-01 | 3.00E-01 | Conventional |
| **Day:TIS** | -5.32E-03 | 8.38E-03 | -6.36E-01 | 6.64E-03 | -8.02E-01 | 2.11E-01 | Conventional |
| **Intercept** | 3.72E-01 | 7.89E-02 | 4.71E+00 | 1.11E-01 | 3.36E+00 | 3.90E-04 | Presenting |
| **Day** | 1.67E-03 | 5.33E-03 | 3.13E-01 | 7.35E-03 | 2.27E-01 | 4.10E-01 | Presenting |
| **EV** | -3.11E-01 | 1.12E-01 | -2.79E+00 | 1.31E-01 | -2.38E+00 | 8.67E-03 | Presenting |
| **TIS** | -4.85E-01 | 1.12E-01 | -4.35E+00 | 1.26E-01 | -3.84E+00 | 6.14E-05 | Presenting |
| **Day:EV** | 2.04E-02 | 7.53E-03 | 2.71E+00 | 8.65E-03 | 2.36E+00 | 9.16E-03 | Presenting |
| **Day:TIS** | 2.60E-02 | 7.53E-03 | 3.45E+00 | 8.49E-03 | 3.06E+00 | 1.11E-03 | Presenting |
| **Intercept** | 1.03E-01 | 6.89E-02 | 1.50E+00 | 5.62E-02 | 1.84E+00 | 3.30E-02 | Proliferating |
| **Day** | -3.11E-03 | 4.61E-03 | -6.75E-01 | 3.57E-03 | -8.72E-01 | 1.92E-01 | Proliferating |
| **EV** | 3.30E-01 | 9.79E-02 | 3.37E+00 | 1.44E-01 | 2.29E+00 | 1.09E-02 | Proliferating |
| **TIS** | 3.72E-01 | 1.04E-01 | 3.56E+00 | 7.69E-02 | 4.84E+00 | 6.62E-07 | Proliferating |
| **Day:EV** | -2.22E-02 | 6.69E-03 | -3.32E+00 | 9.11E-03 | -2.44E+00 | 7.38E-03 | Proliferating |
| **Day:TIS** | -2.31E-02 | 7.29E-03 | -3.16E+00 | 5.04E-03 | -4.58E+00 | 2.32E-06 | Proliferating |
| **Intercept** | 1.17E-01 | 5.51E-02 | 2.12E+00 | 5.96E-02 | 1.96E+00 | 2.48E-02 | Immunosuppressive |
| **Day** | -4.61E-03 | 3.79E-03 | -1.22E+00 | 3.86E-03 | -1.19E+00 | 1.17E-01 | Immunosuppressive |
| **EV** | 9.24E-02 | 8.21E-02 | 1.13E+00 | 7.87E-02 | 1.17E+00 | 1.20E-01 | Immunosuppressive |
| **TIS** | -1.06E+00 | 3.25E-01 | -3.25E+00 | 3.57E-01 | -2.96E+00 | 1.54E-03 | Immunosuppressive |
| **Day:EV** | -5.73E-03 | 5.56E-03 | -1.03E+00 | 4.93E-03 | -1.16E+00 | 1.23E-01 | Immunosuppressive |
| **Day:TIS** | 1.84E-01 | 4.54E-02 | 4.04E+00 | 4.86E-02 | 3.78E+00 | 7.82E-05 | Immunosuppressive |
| **Day-sq:TIS** | -6.67E-03 | 1.56E-03 | -4.28E+00 | 1.61E-03 | -4.15E+00 | 1.67E-05 | Immunosuppressive |

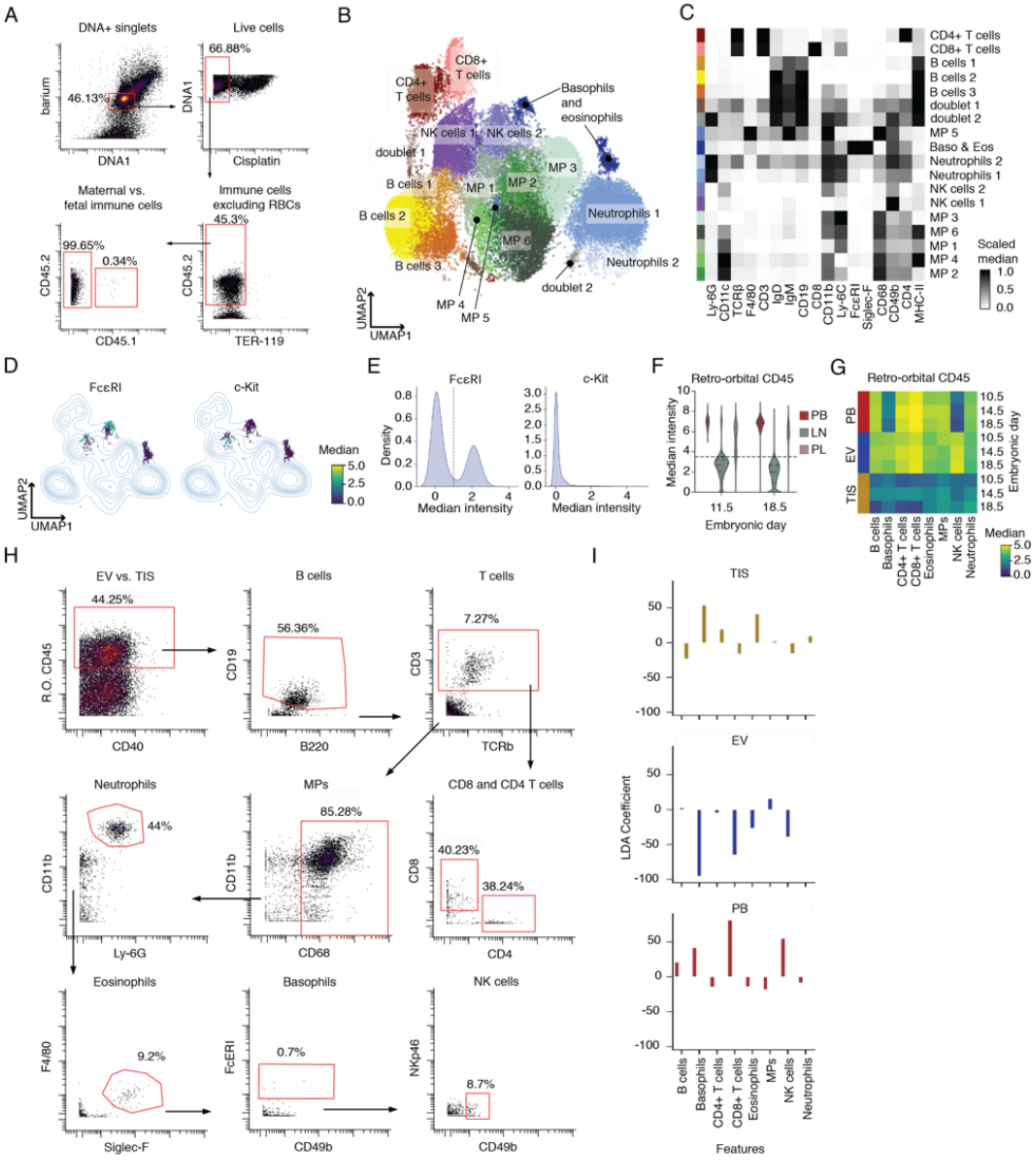
A
DNA+ singlets
barium
46.13%
DNA1
Live cells
66.88%
DNA1
Cisplatin
Maternal vs. fetal immune cells
99.65%
0.34%
CD45.2
CD45.1
Immune cells excluding RBCs
45.3%
CD45.2
TER-119
B
CD4+ T cells
CD8+ T cells
Basophils and eosinophils
NK cells 1
NK cells 2
doublet 1
B cells 1
B cells 2
B cells 3
MP 1
MP 2
MP 3
MP 4
MP 5
MP 6
Neutrophils 1
Neutrophils 2
doublet 2
UMAP2
UMAP1
C
CD4+ T cells
CD8+ T cells
B cells 1
B cells 2
B cells 3
doublet 1
doublet 2
MP 5
Baso & Eos
Neutrophils 2
Neutrophils 1
NK cells 2
NK cells 1
MP 3
MP 6
MP 1
MP 4
MP 2
Ly-6G
CD11c
TCRβ
F4/80
CD3
IgD
IgM
CD19
CD8
CD11b
Ly-6C
FcεRI
Siglec-F
CD68
CD49b
CD4
MHC-II
Scaled median
1.0
0.5
0.0
D
FcεRI
c-Kit
Median
5.0
2.5
0.0
UMAP2
UMAP1
E
FcεRI
c-Kit
Density
Median intensity
F
Retro-orbital CD45
Median intensity
PB
LN
PL
11.5
18.5
Embryonic day
G
Retro-orbital CD45
PB
EV
TIS
10.5
14.5
18.5
Embryonic day
B cells
Basophils
CD4+ T cells
CD8+ T cells
Eosinophils
MPs
NK cells
Neutrophils
Median
5.0
2.5
0.0
H
EV vs. TIS
44.25%
R.O. CD45
CD40
B cells
56.36%
CD19
B220
T cells
7.27%
CD3
TCRb
Neutrophils
44%
CD11b
Ly-6G
MPs
85.28%
CD11b
CD68
CD8 and CD4 T cells
40.23%
38.24%
CD8
CD4
Eosinophils
9.2%
F4/80
Siglec-F
Basophils
0.7%
FcERI
CD49b
NK cells
8.7%
NKp46
CD49b
I
TIS
EV
PB
LDA Coefficient
B cells
Basophils
CD4+ T cells
CD8+ T cells
Eosinophils
MPs
NK cells
Neutrophils
Features

**Figure S1. Maternal immune clusters identified by Leiden, determination of intravascular and extravascular compartments, and gating strategy.** Related to Figure 1. (A) Mass cytometry gating strategy to isolate maternal and fetal immune cells. (B) Leiden clusters overlaid on UMAP graph of maternal immune cells. Leiden clusters were named based on their phenotypic marker expression. (C) Scaled median expression of protein markers used to generate UMAP graph and Leiden clusters are shown across all Leiden clusters identified. (D) UMAP indicating arcsinh transformed cellular median intensity of FcεRI and c-Kit in the Leiden cluster identified as “Basophils and Eosinophils”. (E) Kernel density plots of arcsinh transformed FcεRI and c-Kit medians in the Leiden cluster identified as “Basophils and Eosinophils”. FcεRI plot includes the threshold set to distinguish basophils from eosinophils. (F) Arcsinh transformed single cell median expression of retro-orbitally injected CD45 (R.O. CD45) in matched peripheral blood (PB), lumbar lymph nodes (LN), and placenta (PL). Plot includes the threshold set to distinguish EV vs. TIS compartments in PL. (G) Arcsinh transformed median expression of R.O. CD45 across metaclustered maternal immune cell types. Cell types were split by organ, peripheral blood and placenta. (H) Gating strategy to define EV vs. TIS in placenta E12.5 sample, along with the maternal immune cell types identified. (I) Linear discriminant analysis (LDA) coefficients indicating the weight each feature (cell type) was given to identify each class (compartments TIS, EV, and PB).

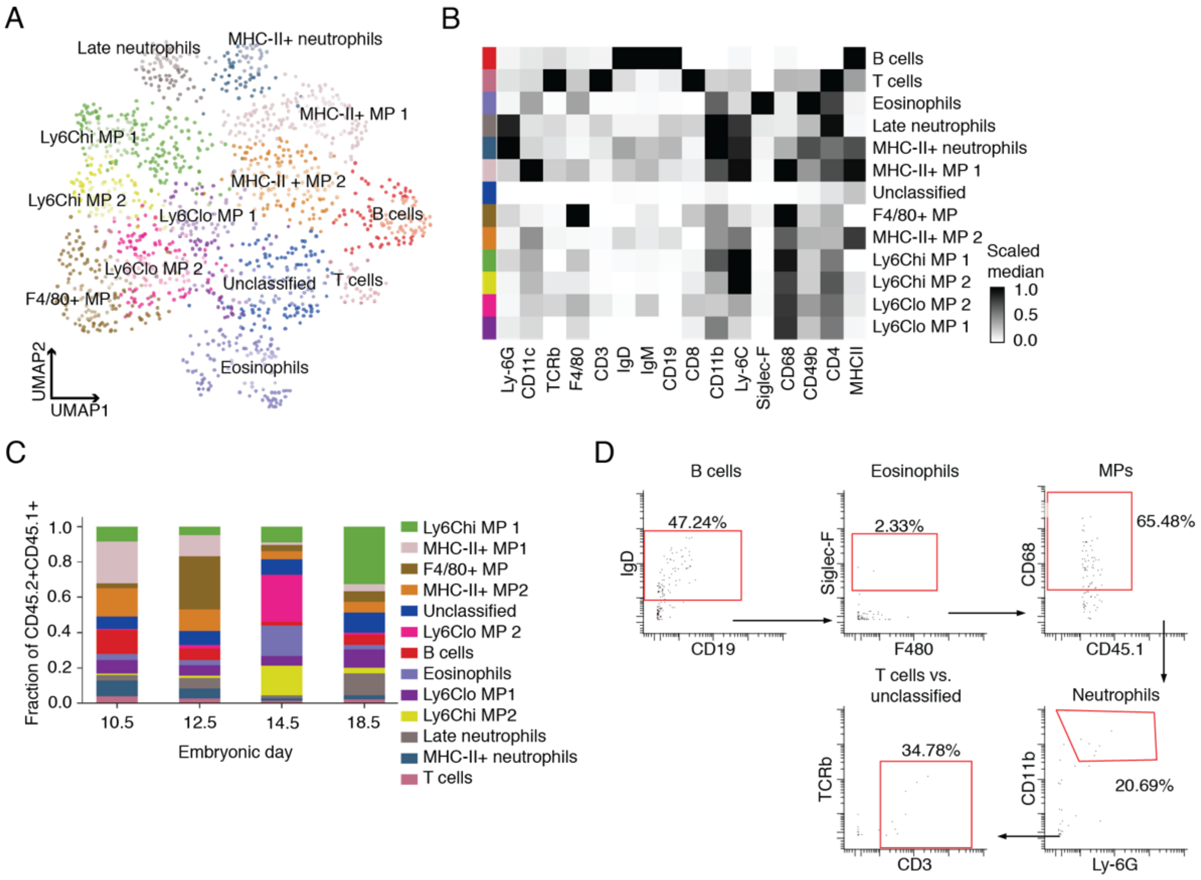


**Figure S2. Fetal immune clusters identified by Leiden and their gating strategy.** Related to Figure 2. (A) Leiden clusters overlaid on UMAP graph of fetal immune cells. Leiden clusters were named based on their phenotypic marker expression. (B) Scaled median expression of protein markers used to generate UMAP graph. Proteins are shown across all Leiden clusters identified. (C) Fraction of fetal immune cells at E10.5, 12.5, 14.5, and 18.5 across Leiden clusters. (D) Gating strategy to identify fetal immune cell types.

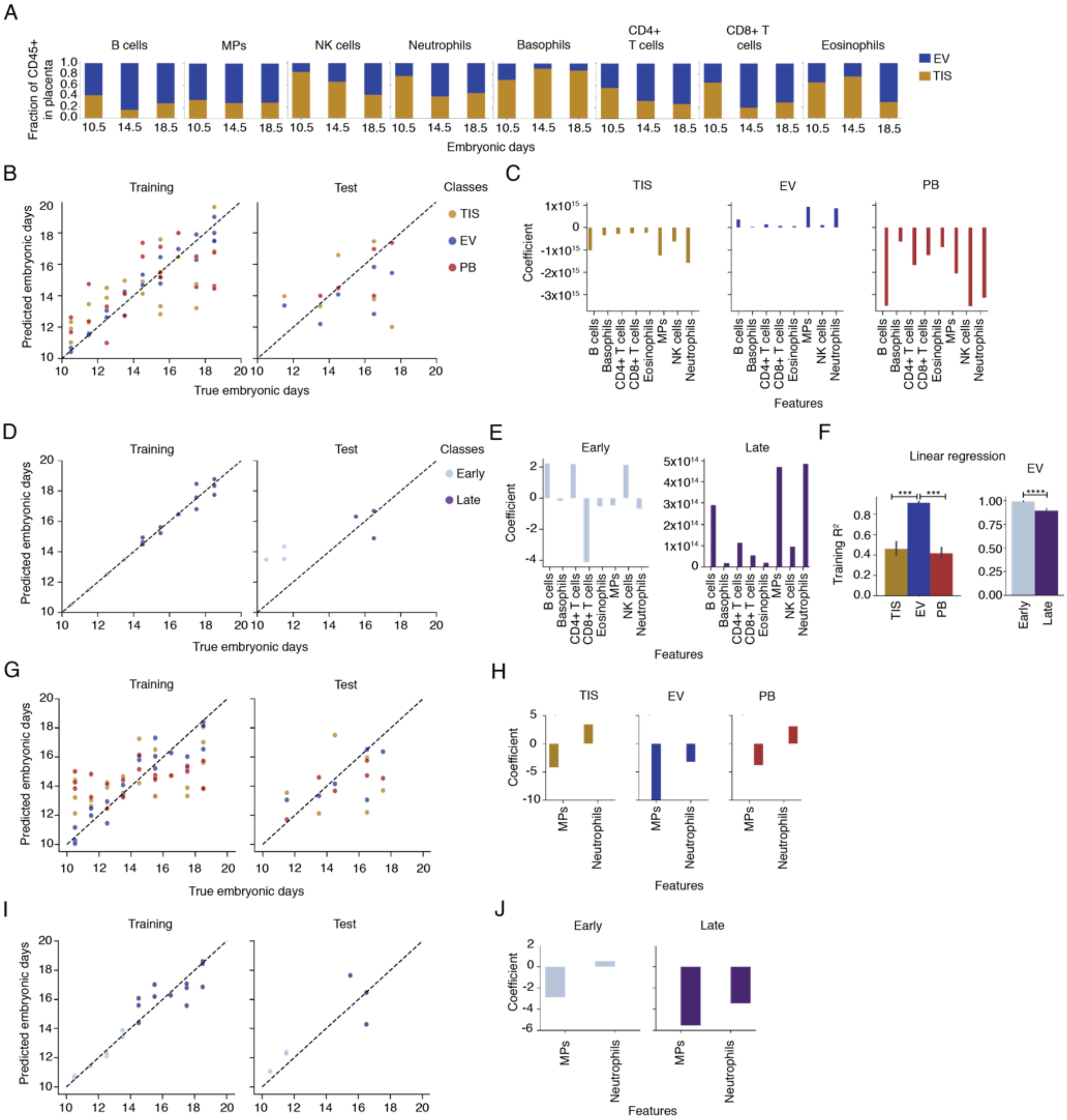
A
Fraction of CD45+ in placenta
B cells
MPs
NK cells
Neutrophils
Basophils
CD4+ T cells
CD8+ T cells
Eosinophils
EV
TIS
Embryonic days
B
Training
Test
Classes
TIS
EV
PB
Predicted embryonic days
True embryonic days
C
TIS
EV
PB
Coefficient
Features
D
Training
Test
Classes
Early
Late
Predicted embryonic days
True embryonic days
E
Early
Late
Coefficient
Features
F
Linear regression
EV
Training R²
G
Training
Test
Predicted embryonic days
True embryonic days
H
TIS
EV
PB
Coefficient
Features
I
Training
Test
Predicted embryonic days
True embryonic days
J
Early
Late
Coefficient
Features

**Figure S3. Frequencies of maternal immune cell subsets show predictive dynamics throughout gestation.** Related to Figure 3. (A) Fraction of maternal immune cells in the placenta extravascular (TIS) and intravascular (EV) compartments across embryonic days 10.5, 14.5, and 18.5. (B) Linear regression results by maternal compartment (classes) based on maternal immune cell fractions out of all immune cells in the given compartment (features). The training set is made up of previously seen data, and the test set is made up of data that has not been seen by the linear regression algorithm. (C) Linear regression coefficients for features, in this case cell type, within the specified class, or maternal compartment. (D) Cell fractions of EV compartment were split into early (E10.5 - 13.5) and late (E14.5 – 18.5) and linear regression was run independently for the early and late periods. Linear regression results on the training and test sets based on maternal immune cell fractions (features). (E) Linear regression coefficients for features, in this case cell type, within the specified class, or EV embryonic period. (F) Training accuracy as determined by $R^2$ of linear regression across each compartment and EV period based on cell fractions across embryonic days. ***$p \leq 0.001$, ****$p \leq 0.0001$ (one-way ANOVA for comparing compartments, unpaired t test for early and late stages). (G) Linear regression results by maternal compartment (classes) based on maternal immune cell fractions of mononuclear phagocytes and neutrophils in the given compartment (features). (H) Linear regression coefficients for features, in this case cell type, within the specified class, or maternal compartment. (I) Cell fractions of mononuclear phagocytes and neutrophils EV compartment were split into early and late and linear regression was run independently for the early and late periods. (J) Linear regression coefficients for features, in this case cell type, within the specified class, or EV embryonic period.

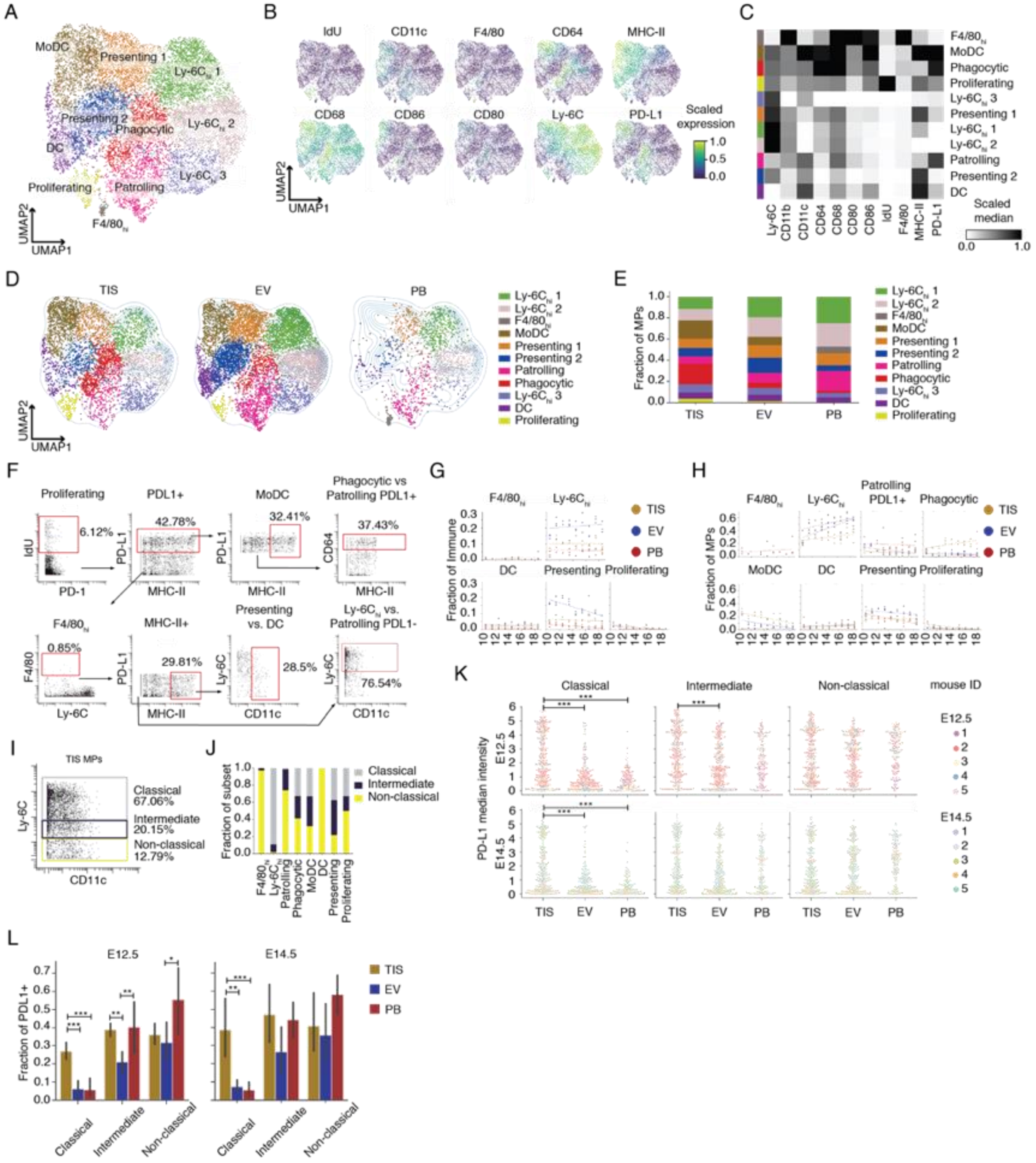

A
B
C
D
E
F
G
H
I
J
K
L
UMAP1
UMAP2
Scaled expression
Scaled median
TIS
EV
PB
Fraction of MPs
Fraction of Immune
Fraction of subset
Fraction of PDL1+
PD-L1 median intensity
E12.5
E14.5
Classical
Intermediate
Non-classical
mouse ID
TIS MPs

**Figure S4. Maternal mononuclear phagocyte clusters identified by Leiden, their gating strategy, and canonical monocyte subset phenotypes.** Related to Figure 4. (A) Leiden clusters overlaid on composite UMAP graph of maternal mononuclear phagocytes (MPs) in placenta and peripheral blood across E10.5 to 18.5. Leiden clusters were named based on their phenotypic marker expression. (B) Scaled cellular median intensity of MP phenotypic markers. (C) Scaled median expression of protein markers are shown across all Leiden clusters identified. (D) UMAP plots showing distribution of MP clusters identified by Leiden across compartments. (E) Fraction of MP clusters out of total MPs in each compartment. Embryonic days were aggregated. (F) Gating strategy to identify MP subsets. (G) Fraction of PD-L1 negative MP subsets out of maternal immune cells were fitted with GEE using a linear model comparing TIS, EV, and maternal PB from E10.5 to 18.5. (H) GEE was used to fit a linear model to cell fractions of MP subsets out of all MPs comparing TIS, EV, and PB from E10.5 to 18.5. (I) Gating strategy to identify canonical MP subsets, classical, intermediate, and non-classical. (J) Fraction of canonical MP classification for each MP subset. (K) Single cell arcsinh median intensity of PD-L1 in canonical MP subsets across compartments and at E12.5 and 14.5. To clearly show single cell distribution, samples were capped at 30 cells per mouse. (L) PD-L1 positive fraction of each canonical MP subsets across compartments and at E12.5 and 14.5. For (K) and (L), significance is shown as *$p \leq 0.05$, **$p \leq 0.01$, ***$p \leq 0.001$ (one-way ANOVA per cell type).

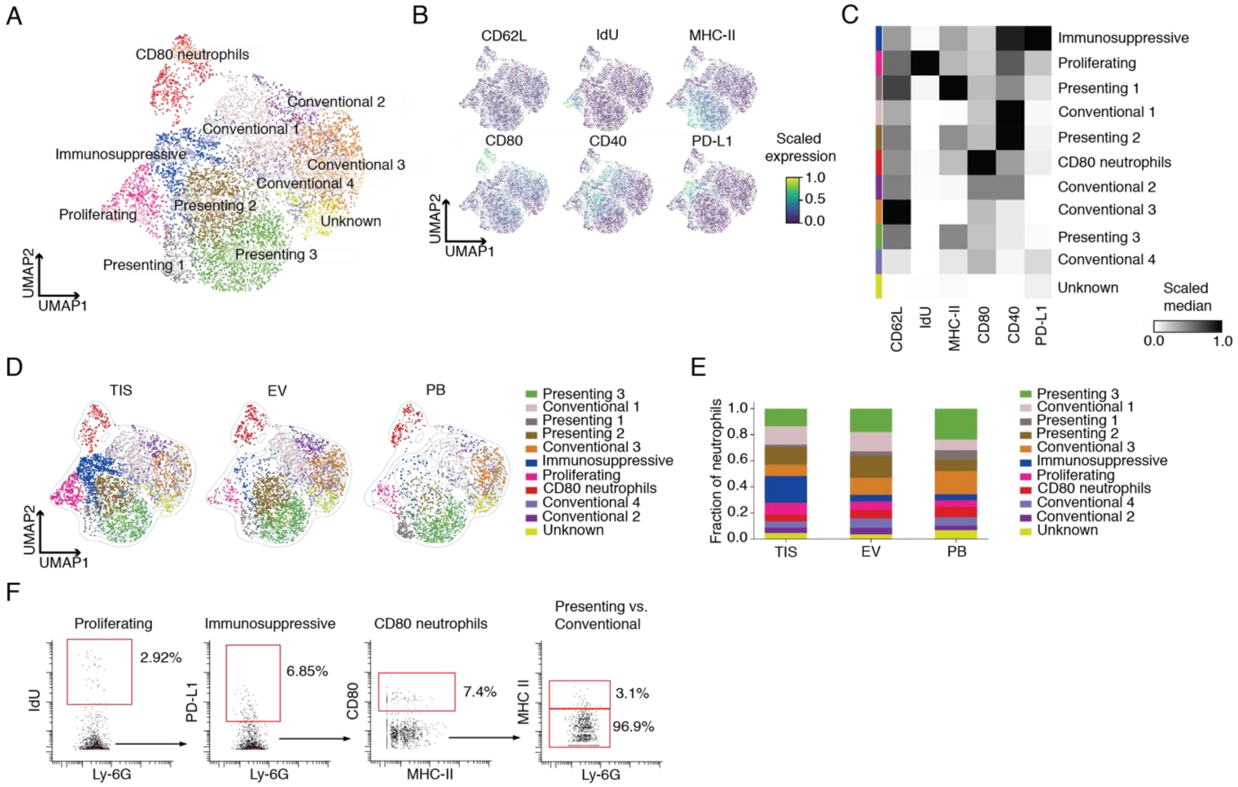


**Figure S5**. **Maternal neutrophil clusters identified by Leiden and their gating strategy.** Related to Figure 5. (A) Leiden clusters overlaid on composite UMAP graph of maternal neutrophils in placenta and peripheral blood across E10.5 to 18.5. Leiden clusters were named based on their phenotypic marker expression. (B) Scaled cellular median intensity of neutrophil phenotypic markers. (C) Scaled median expression of protein markers are shown across all Leiden clusters identified. (D) UMAP plots showing distribution of neutrophil clusters identified by Leiden across compartments. (E) Fraction of neutrophil clusters out of total neutrophils in each compartment. Embryonic days were aggregated. (F) Gating strategy to identify neutrophil subsets.

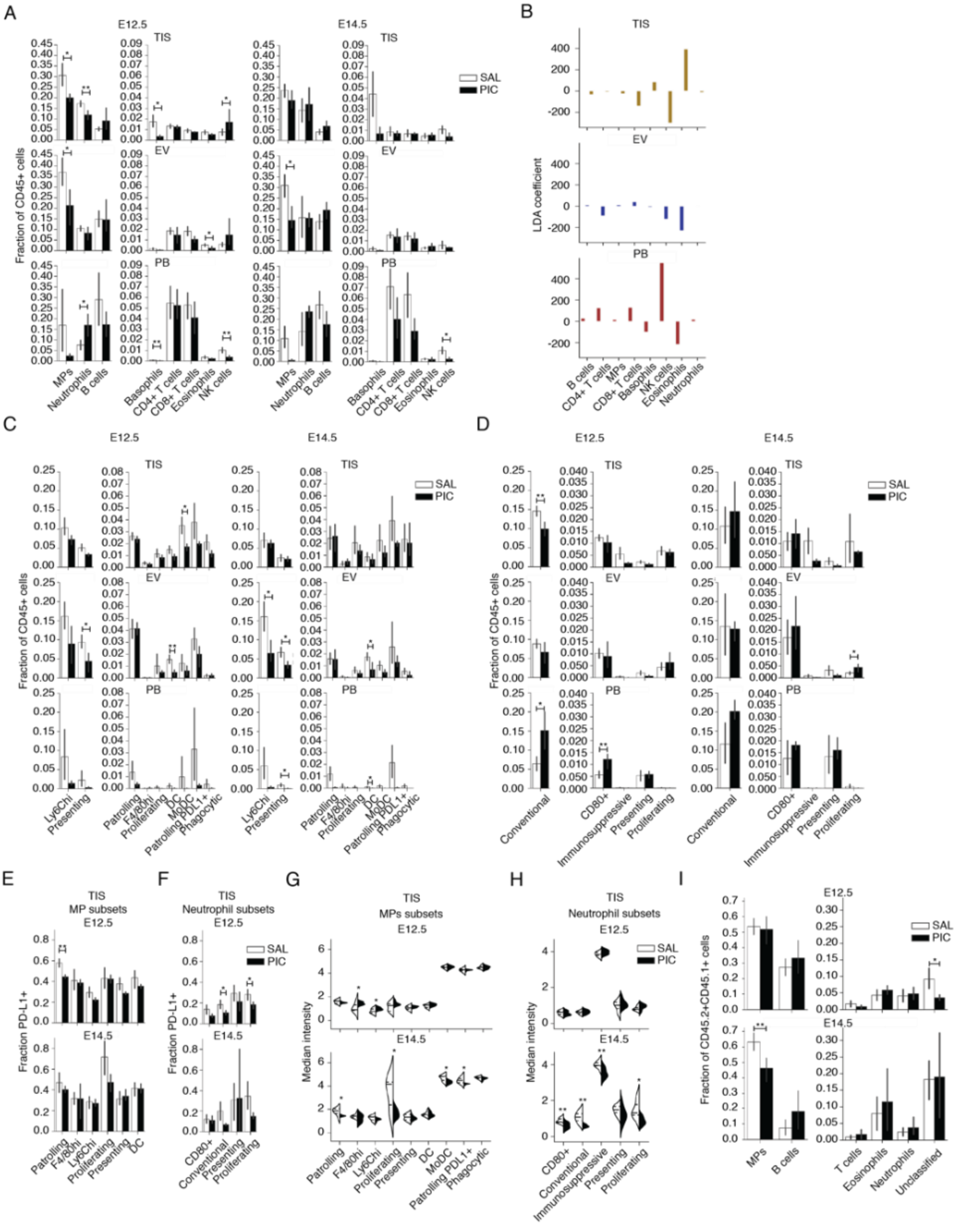

A
E12.5
E14.5
TIS
EV
PB
SAL
PIC
Fraction of CD45+ cells
MPs
Neutrophils
B cells
Basophils
CD4+ T cells
CD8+ T cells
Eosinophils
NK cells
B
LDA coefficient
C
Ly6Chi
Presenting
Patrolling
F4/80hi
Proliferating
DC
MoDC
Patrolling PDL1+
Phagocytic
D
Conventional
CD80+
Immunosuppressive
E
MP subsets
Fraction PD-L1+
Ly6Chi
F
Neutrophil subsets
G
MPs subsets
Median intensity
H
I
Fraction of CD45.2+CD45.1+ cells
T cells
Eosinophils
Unclassified

**Figure S6**. **Maternal and fetal immune responses to maternally, systemically administered Poly(I:C) at mid gestation.** Related to Figure 6. (A) For E12.5 and 14.5, maternal immune cell type fractions across compartments when challenged with saline (SAL) or Poly(I:C) (PIC).. (B) Linear discriminant analysis (LDA) coefficients for cell type (features), for each compartment (class). LDA was trained on baseline data comprised of E12.5 and 14.5. (C) For E12.5 and 14.5, fractions of maternal MP subsets across compartments when challenged with SAL or PIC. (D) For E12.5 and 14.5, fractions of maternal neutrophil subsets across compartments when challenged with SAL or PIC. (E) PD-L1 positive fractions of PD-L1 low maternal MP subsets at E12.5 and 14.5, in TIS compartment when challenged with SAL or PIC. (F) PD-L1 positive fractions of PD-L1 low maternal neutrophil subsets at E12.5 and 14.5, in TIS compartment when challenged with SAL or PIC. (G) Arcsinh transformed median intensity of PD-L1 for MP subsets in TIS at E12.5 and 14.5 upon SAL or PIC challenge. (H) Median intensity of PD-L1 for neutrophil subsets in TIS at E12.5 and 14.5 upon SAL or PIC challenge. (I) For E12.5 and 14.5, fetal immune cell fraction was compared for immune challenge with SAL or PIC. For (A) and (C) through (I), significance is shown as *$p \leq 0.05$, **$p \leq 0.01$, ***$p \leq 0.001$ (unpaired t test).

### TRANSPARENT METHODS

##### *Animals*

All mice were housed in an animal facility that is accredited by the Association for Assessment and Accreditation of Laboratory Animal Care International and maintained in specific pathogen-free conditions. Animal studies were conducted in accordance with National Institutes of Health guidelines for the humane use of animals and reviewed and approved by the Stanford Institutional Animal Care and Use Committee. Wild-type female and male C57BL/6J mice between 6-8 weeks old were purchased from The Jackson Laboratory and housed at our facility. To differentiate between maternal and fetal cells, male C57BL/6 CD45.1 (B6.SJL-Ptprca Pepcb/BoyJ) mice between 7-8 weeks old were also purchased from The Jackson Laboratory. Animals were housed under standard 12-hour light/dark cycles and fed with standard chow.

##### *Timed Pregnancies and Treatments*

Timed pregnancies were generated by housing one or two naturally cycling females with a single male overnight. Mice were separated early the following morning and mating was assessed by the appearance of a copulation plug. The day of plug detection was classified as E0.5. For baseline studies, pregnant mice at embryonic days 10.5, 11.5, 12.5, 13.5, 14.5, 15.5, 16.5, 17.5, and 18.5 were injected intraperitoneally (i.p.) with saline 2-3 hours before sacrifice. For perturbation studies, at E12.5 or 14.5, pregnant mice were treated with Poly(I:C) (Invivogen, tlrl-picw). Poly(I:C) was dosed at 18.5 mg/kg body weight, prepared at a concentration of 5 mg/mL in saline and injected i.p. 2-3 hours before sacrifice. To assess proliferative activity, pregnant mice received one intraperitoneal injection of Iododeoxyuridine (IdU; Sigma-Aldrich, #I7125). IdU was dissolved in saline after bringing pH to 10 with sodium hydroxide and incubating on a 37°C shaker. Once dissolved, pH was brought to 8.5 using hydrochloric acid and solution was filtered before injection. IdU was dosed at 100 mg/kg of body weight at a concentration of 10 mg/mL 2-3 hours before sacrifice.

##### *Detection of Endovascular Immune Cells*

Mice were injected retro-orbitally at least 2 min before sacrifice with up to 5 µg biotin rat anti-mouse CD45 Ab (clone 30-F11; BioLegend, #103104) in 60 µl saline. We analyzed samples by mass cytometry to determine successful anti-CD45 labeling in peripheral blood and its absence in matched cells from the lumbar lymph nodes (Figure S1F). Mice that showed significant Ab leakage into lymph nodes were excluded from downstream analysis.

##### *Tissue Preparation and Cell Isolation*

After terminal anesthesia by ketamine and xylazine, peripheral blood was collected via cardiac puncture and transferred into K2EDTA evacuated blood collection tubes (Fisher Scientific, #02-683-99A). Cells from the peripheral blood were subjected to RBC Lysis Buffer (BioLegend, #420301) to remove red blood cells. Tissue was collected from each fetus for sex genotyping and stored at -20°C. Placentas (with decidua) were collected and minced with scissors in cold Accutase (Sigma Aldrich, #SCR005) before transferring to a 37°C incubator shaker for enzymatic digestion as previously described (Arenas-Hernandez et al., 2015). Following digestion, samples were centrifuged at 1200 rpm for 2 min at RT. Placentas were not dissected into fetal and decidual (maternal) portions. Placentas were processed whole to reduce variability, preserve the capacity to compare samples across gestation, and maintain sample integrity. Complete decidual removal from the fetal spongiotrophoblast and trophoblast giant cells, while never guaranteed, was only effectively possible on a subset of embryonic days. Inconsistent decidual removal would prevent cross-gestational analysis of the placenta. Additionally, decidual removal required unavoidable physical pressure on the placenta, which resulted in endovascular leakage. Any leakage of this sort would prevent us from systematically profiling the endovascular compartment of the placenta. Samples were filtered through a 70 micron cell strainer (Falcon). All single-cell suspensions were quenched and washed with FACS buffer (PBS with 10% FCS and 5 mm EDTA) at 4°C. To label non-viable cells, all cells were resuspended at a 1:1 ratio with PBS with 5 mm EDTA and 100 µM cisplatin (Sigma Aldrich, #P4394) for 1 min before quenching at a 1:1 ratio in FACS buffer. Cells were centrifuged at 1200 rpm for 5 min at 4°C and resuspended in FACS buffer and fixed for up to 1 hour at RT using the FOXP3 Transcription Factor Staining Buffer Set (eBioscience, #00-5523-00) at a cell density of 1 million cells per 500 µl final volume. Cells were then resuspended in FACS buffer and centrifuged at 1500 rpm for 5 min at 4°C. Samples were kept at 4°C during all steps of tissue harvest and cell isolation except enzymatic digestion, viability staining, and fixation. Cells were stored at -80°C until all samples were ready for staining.

##### *Sex Genotyping*

DNA was extracted from fetal body tissue for sex genotyping (Sigma Aldrich, #XNAT). Primers used for sex genotyping PCR target the X-chromosome-specific gene Jarid1c and the Y-chromosome-specific gene Jarid1d (Forward primer: 5-CTGAAGCTTTTGGCTTTGAG-3; Reverse primer: 5-CCGCTGCCAAATTCTTTGC-3). Female samples exhibit a single band at 331 bp, whereas male samples have two bands at 302 and 331.

##### *Mass Cytometry Antibody Conjugation*

Antibody conjugation was performed as previously described (Hartmann et al., 2019). Briefly, metal-isotope labeled antibodies were conjugated using the MaxPar X8 Antibody Labeling kit according to the manufacturer's protocol (Fluidigm) or were purchased pre-conjugated (Fluidigm). To validate conjugation, the absorbance of the conjugated antibody was measured at 280 nm and the concentration was calculated, often resulting in over 60% recovery of antibody. Antibodies were titrated to determine the optimal staining concentration using primary mouse cells and/or mouse cell lines. For long-term storage at 4°C, antibodies were diluted in Antibody Stabilizer solution (Candor Bioscience GmbH, #131-050) with 0.02% NaN3 (Merck Chemicals, #106688) at 0.2 mg/mL. All mass cytometry antibodies and concentrations used in these studies can be found in Table S1.

##### *Mass Cytometry Sample Processing and Data Acquisition*

Placentas were pooled separately by sex for each litter. Mass-tag cell barcoding was employed as previously described[3] to pool samples for more efficient processing and measurement. Briefly, each sample was labeled with distinct combinations of six stable Pd isotopes in PBS with 0.02% saponin. Barcoded samples were washed with cell staining media (CSM; PBS with 0.5% BSA and 0.02% NaN3; Sigma Aldrich) and pooled into a single 5 mL round-bottom polystyrene test tube (Corning) for surface staining. Barcoded samples were suspended in TruStain FcX (BioLegend, #101320) to prevent non-specific antibody binding and incubated on ice for 10 min prior to staining. Surface staining was performed in CSM in 500 µL total volume for 30 min at RT. Cells were washed in CSM and fixed (eBioscience, #00-5523-00) for 10 min at RT. Cells were centrifuged at 1600 rpm for 5 min at 4°C and supernatant was aspirated after all washes. Cells were washed once in CSM and once in permeabilization buffer (eBioscience, #00-5523-00). Cells were stained with intracellular antibodies in permeabilization buffer in 500 µL total volume for 30 min at RT. Cells were washed in CSM and stained with 1 mL DNA intercalation solution (1.6% PFA in low barium PBS with 0.02% saponin and 0.5 μM Cell-ID Intercalator-Ir; Fluidigm, #201192) overnight at 4°C or until data acquisition, not exceeding 7 days. Before data acquisition, samples were washed once in CSM and twice in ddH2O. All samples were resuspended in 1x EQ Four Element Calibration Bead solution (Fluidigm, #201078) with ddH2O at 1-2x$10^6$ cells/mL and filtered through a cell strainer capped test tube (Falcon, #352235) before being injected into a CyTOF2+ mass cytometer (Fluidigm) using the Super Sampler injection system (Victorian Airship and Scientific Apparatus).

***Mass Cytometry Data Processing***

After cell acquisition, FCS files for each sample were bead normalized and concatenated with the ParkerICI/premessa package in R (https://rdrr.io/github/ParkerICI/premessa/f/README.md). FCS files obtained from barcoded plates were then deconvoluted with the Single Cell Debarcoder application developed by Zunder et al., 2015 (Zunder et al., 2015). To correct for technical variation between CyTOF runs, we quantile normalized protein expression with the Cydar package in R (https://www.bioconductor.org/packages/release/bioc/html/cydar.html). Each barcode plate included a splenocyte sample to which we normalized all samples across plates. These FCS files were then uploaded to Cell Engine for gating (Figure S1A). All parameters except for time and cell length were displayed with an arcsinh cofactor 5 transformation. Events positive for intercalator-Ir were selected as having high DNA content. Cisplatin was then used to discriminate between live and dead cells. Staining with TER119 allowed exclusion of red blood cells from proceeding gates. Cells were then either gated for their expression of CD45.2+ single positive, deemed maternal immune, or CD45.2+CD45.1+ double positive, deemed fetal immune (Figure S1A).

We established whether placental cells were in tissue (TIS) or within the endovasculature (EV) by setting a threshold based on the expression of retro-orbitally (R.O.) injected CD45-biotin in peripheral blood (PB) and lumbar lymph nodes (LN) (Figure S1F). With the arcsinh cofactor 5 transformation of cellular medians, we set the threshold to equal 3.5. In the placenta samples, any cells with median intensity equal to or higher than 3.5 were considered to be in EV. Any cells with R.O. CD45 that fell below the 3.5 threshold were considered to be in TIS. When we visualize the median intensity of R.O. CD45 across entire samples, we see a distinction in the expression of R.O. CD45 in EV vs. TIS (Figure S1G). We confirmed detection of EV immune cells by traditional gating as well (Figure S1H).

The maternal and fetal immune cell subsets identified in Figures 1 and 2 through dimensionality reduction and clustering were then confirmed using traditional gating methodology on Cell Engine as seen in Figures S1H and S2D. We back-gated to ensure cells were not present in multiple gates. Furthermore, we identified the maternal mononuclear phagocyte (Figure 4A) and neutrophil (Figure 5A) subsets via traditional gating in Figures S4F and S5F, respectively. To ensure every cell was counted, we gated in a hierarchical manner as shown (Figures S4F and S5F).

Canonical maternal MP subsets were determined by their Ly-6C expression. Based on statistics of Ly-6C MP expression in peripheral blood, MPs were considered “classical” if their arcsinh cofactor 5 median expression was equal or higher than 4.5. Fiftieth percentile was 4.9, mean was 4.1, and standard deviation was equal to 2. “Intermediate” MPs were those with Ly-6C expression equal or higher than 3 but lower than 4.5 ($25^{th}$ percentile was 2.7). Lastly, “non-classical” MPs had Ly-6C median intensity lower than 3.

***Data Analysis***

**Dimensionality reduction and clustering**

We used Scanpy's Python based implementation (Wolf et al., 2018) to carry out dimensionality reduction via UMAP and clustering with the Leiden algorithm. These were carried out separately for maternal and fetal immune cells. Protein median intensities were first transformed with an inverse hyperbolic sine (arcsinh) with a cofactor of 5. We computed a UMAP neighborhood graph from baseline maternal immune data by randomly subsampling up to 500 single positive CD45.2+ cells from each mouse organ from embryonic day 10.5 to 18.5 and restricting local neighbors to 20. UMAP was based on the expression of the following lineage markers: Ly-6G, CD11c, TCRb, F4/80, CD3, IgD, IgM, CD19, CD8, CD11b, Ly-6C, FcεRI, Siglec-F, CD68, CD49b, CD4, and MHC-II. These same lineage markers were used to carry out Leiden clustering (Figure S1B).

Leiden produced 16 clusters, including 2 clusters that were excluded from downstream analysis due their doublet inclusion and broad expression of all lineage markers. With the remaining 14 clusters, we hierarchically clustered 2 B cell clusters, 2 neutrophil clusters, and 2 NK cell clusters into their broader cell types. Leiden also produced 5 mononuclear phagocyte (MP) clusters which had a range in expression of Ly-6C, MHC II, and CD11b. These clusters likely include macrophages (Mac), dendritic cells (DCs), monocytes, and monocytes differentiating into Mac or DCs. We decided to hierarchically cluster these cells because their over-clustering could be due to their low expression of other lineage markers, resulting in non-specific cellular distributions in the UMAP (Figure S1B-C). We acknowledge that heterogeneity also contributed to the generation of multiple clusters per cell type, but analyzing heterogeneity is best done by applying additional clustering markers that are relevant to and sufficiently expressed by these cells, as we did for MPs (Figure 4) and neutrophils (Figure 5). Finally, we identified one Leiden cluster that was composed of both eosinophils and basophils and spatially separated in the UMAP graph (Figure S1B, S1D). We set a threshold of arcsinh cofactor 5 transformed FcεRI median equal to 1 to split this single cluster into two distinct populations (Figure S1E). FcεRI is specific to basophils. Furthermore, we found very low levels of c-Kit in the "Basophils and Eosinophils" Leiden cluster (Figure S1D), suggesting the inclusion of mast cells. The c-Kit positive population was found to overlap with the location of NK cells in the UMAP graph. In the kernel density plot of c-Kit (Figure S1E), the levels of c-Kit positive cells were overwhelmed by the vastly c-Kit negative "Basophil and Eosinophil" population. Because we had such a low number of these mast cells, we decided to keep them in the heterogenous eosinophil population.

We computed a UMAP neighborhood graph from baseline fetal immune data by subsampling up to 150 CD45.2+CD45.1+ cells from placentas on embryonic days 10.5, 12.5, 14.5, and 18.5, then applied the same Leiden setting used for the maternal immune cell analysis. Applying Leiden resulted in 13 clusters (Figure S2A). Seven of these clusters were classified as MPs with differential MHC-II, F4/80, and Ly-6C expression (Figure S2B), and were grouped into a single cluster. Similarly, two clusters were identified as neutrophils with differential MHC-II expression (Figure S2B) and were grouped into a single cluster. Finally, there was one cluster left unassigned because its marker expression was low for all lineage markers tested.

To further analyze maternal MP heterogeneity, we isolated maternal MPs and applied UMAP and Leiden using biologically relevant markers expressed in MPs: CD11c, F4/80, CD64, CD68, CD86, CD80, MHC-II, and cellularly incorporated IdU to track cell proliferation (Figure 4). The UMAP neighborhood graph was again restricted to 20 neighbors. Applying Leiden resulted in 11 MP clusters (Figure S4A). We found 3 Leiden clusters that were nearly identical based on their median protein expression (Figure S4C), so we grouped them into a single "Ly-6C$_{hi}$" subset. Additionally, two MHC-II expressing clusters with negative CD11c were grouped into the "Presenting" subset. We removed a F4/80 high expressing cluster from downstream analysis because it spatially overlapped with several other clusters on the UMAP graph (Figure S4A and S4D). The "F4/80$_{hi}$" cluster was only found in the peripheral blood.

To further analyze maternal neutrophil heterogeneity, we isolated maternal neutrophils and applied UMAP and Leiden using CD62L, MHC-II, CD80, CD40, PD-L1, and incorporated IdU to track cell proliferation (Figure 5). The UMAP neighborhood graph was restricted to 20 neighboring cells. Applying Leiden resulted in 11 clusters, which were then grouped based on the differential expression of CD11b, Ly-6C, Ly-6G, and CD44 in addition to the markers used for clustering (Figure S5). Four Leiden clusters were grouped under "conventional" and 3 under "presenting" neutrophils. One of the clusters Leiden identified was negative for all markers tested, including Ly-6G, so we removed it from further analysis.

We also removed from analysis technical artifacts and sample outliers if we observed metal isotope bleed-through, sample-specific clusters, and anomalous sample-driven clustering.

**Linear Discriminant Analysis**

We dimensionally reduced cell fractions of B cells, basophils, CD4 T cells, CD8 T cells, eosinophils, mononuclear phagocytes, NK cells, and neutrophils across the three compartments analyzed (TIS, EV, and PB) by implementing SciKit Learn's Linear discriminant analysis (LDA). For Figure 1G, the compartments served as the three class labels, and the cell fractions within each compartment were the features. The LDA coefficients can be found in Figure S1I. In Figure 6B, we trained the LDA on saline samples (contour plots) and transformed the input cell frequencies from samples of Poly(I:C)-challenged mice. Poly(I:C) samples were then overlaid as points with their original class label. The LDA coefficients for this analysis be found in Figure S6B.

**Bray-Curtis Index of Dissimilarity**

Beta diversity is a ratio metric used in ecology to measure the degree of difference in species composition across communities or environments. We considered immune cells to be similar to a community of species. We used the scikit-bio.diversity beta subpackage (http://scikit-bio.org/docs/0.2.0/generated/skbio.diversity.beta.html) and Bray-Curtis metric to measure the degree of difference between the three compartments analyzed (TIS, EV, and PB) using the cell abundance of B cells, basophils, CD4 T cells, CD8 T cells, eosinophils, mononuclear phagocytes, NK cells, and neutrophils (Figures 1H and 6C).

**Microarray Data**

Publicly available mouse placenta and decidua microarray data (Knox & Baker) was analyzed for immune and vascular development genes (Figure 3B). The immune gene dataset was obtained from Immport (https://www.immport.org/resources), and the vascular development gene dataset was obtained from JAX (http://www.informatics.jax.org/go/term/GO:0001944). We implemented R to determine differentially expressed genes with the aid of the limma package (https://academic.oup.com/nar/article/43/7/e47/2414268) and focused on genes that changed between E8.5 and 15.5. The full dataset of immune and vascular associated genes we found to be differentially expressed between E8.5 and 15.5 can be found in the supplemental files.

#### Linear Regression

We used scikit-learn's implementation of linear regression to determine if the frequency of B cells, basophils, CD4 T cells, CD8 T cells, eosinophils, mononuclear phagocytes, NK cells, and neutrophils demonstrated temporal organization across the three compartments analyzed (Figure S3B). The features we used were the fractions of the given cell types (calculated out of all immune cells in the compartment) and our target values were embryonic days. Linear regression coefficients are shown in Figure S3C. We used scikit-learn's cross validation feature to determine the Ridge regression score function ($R^2$), allowing us to evaluate our model's embryonic day prediction based on immune cell composition. The $R^2$ when using all cells as features is found in S3F. We additionally applied linear regression only using the frequency of MPs and neutrophils across compartments recapitulating a similar pattern in accuracy scores when comparing compartments (Figure 3E). We carried out linear regression in the endovascular (EV) compartment in early (E10.5 to 13.5) and late (E14.5 to 18.5) periods of gestation using all cell types (Figure S3D). The coefficients of this regression can be found in Figure S3E. The $R^2$ scores using all cell types as features for EV gestational period are shown in Figure S3F, while the score for using only MPs and neutrophils can be found in Figure 3E.

#### Generalized Estimating Equations

To estimate the effect of gestational day across placental compartments (Figures 3, 4, and 5), linear regression coefficient and standard error estimates were calculated using the Generalized Estimating Equations (GEE) framework (Liang and Zeger, 1986). Separate regressions were run depending on cell type, cell function, or protein marker. In each regression, the main effects of gestational day and compartment were included along with the interaction with day and day-squared where appropriate. Using a cluster size of 1 and an independent correlation structure, this approach was equivalent to using heteroskedastic-robust standard errors where the variance of error terms are not identically distributed, but are instead estimated using the squared residual from the individual observations (Eicker, 1967; Huber, 1967; White, 1980).

For comparing average median protein intensity between compartments within a given cell type, the same regression models were run using only the main effects of the EV and TIS compartments without gestational day producing coefficient estimates which compared each compartment against PB. Similar results can be accomplished via a one sided t-test.

All regressions were run in R (v4.1.0) using the "gee" package (v4.13-20). Controlling for heteroskedasticity was justified by visual examination of the residuals plotted against the fitted values in each regression category.

#### Statistics

Statistical tests comparing means of two independent samples were performed with the assistance of the SciPy statistics module using T tests. To calculate p values for two ratio comparisons (Figures 6H and 6I), we applied SciPy's T test for two independent samples from descriptive statistics. To calculate the standard deviation of the ratio of two independent variables (Figures 6H and 6I), we took the square root of the variance, which was calculated using the Taylor Series: $V(X/Y) = E(X_2/Y_2) - [E(X/Y)]_2 = E(X_2)\cdot E[(1/Y)_2] - [E(X)\cdot E(1/Y)]_2$. When indicated, we adjusted p values with the Bonferroni method using the Python module statsmodels (Figure 3F).

One-way analysis of variance (ANOVA) was used when comparing three or more means of independent samples. Pingouin (Vallat, 2018) was used to determine homogeneity of variances, apply classic ANOVA if the groups being compared had equal variances or use the Welch ANOVA for groups with unequal variances. Additionally, the Tukey-HSD post-hoc test was used following a classic ANOVA, and the Games-Howell test was used for samples with unequal variances.

Significant P values are shown as follows on figures: *$p \leq 0.05$, **$p \leq 0.01$, ***$p \leq 0.001$, ****$p \leq 0.0001$.

Figures were created using BioRender (http://www.biorender.com).
